# Notch1 regulates breast cancer stem cell function via a non-canonical cleavage-independent pathway

**DOI:** 10.1101/2020.02.28.970764

**Authors:** Lufei Sui, Suming Wang, Roberto K. Rodriguez, Danielle Sim, Nandita Bhattacharya, Anna L. Blois, Siqi Chen, Sura Aziz, Thorsten Schlaeger, Michael S. Rogers, Diane Bielenberg, Lars A. Akslen, Randolph S. Watnick

## Abstract

Current treatment of triple negative breast cancer patients is hindered by a high incidence of chemoresistance (30-50%). The prevailing theory is that resistance and subsequent recurrence is driven by cancer stem cells. Unfortunately, the functional characterization of cancer stem cells at the molecular level is still incomplete. We show here, that within the canonical breast cancer stem cell population, a subset of cells characterized by high Notch1 expression possesses the tumor-initiating property associated with cancer stem cells. Moreover, the tumor initiating property of these high Notch1-expressing breast cancer stem cells is mediated by a cleavage independent Notch signaling pathway culminating in the repression of SIRT1. Of note, the Notch1-mediated repression of SIRT1 is required not only for tumor initiation, but also for chemoresistance in breast cancer stem cells. Strikingly, inhibition of SIRT1 obviates the requirement for Notch1, marking the first example of conferring cancer stem cell function by inhibiting the activity of a single protein. We also demonstrate that progenitor-like mammary epithelial cells, which possess both luminal and basal properties, are also characterized by high Notch1 expression and repression of SIRT1 via the non-canonical pathway. These findings provide the first functional mechanistic requirements for tumor initiation by breast cancer stem cells and suggest that activation of the non-canonical Notch1 pathway is hardwired into tumor-initiating progenitor cells and thus a prerequisite for tumor initiation.

**Statement of Significance:** We demonstrate that chemoresistant and tumor-initiating properties of breast cancer stem cells are driven by repression of SIRT1 via non-canonical Notch signaling, suggesting a novel therapeutic strategy for triple negative breast cancer.

## Introduction

Treatment of triple negative breast cancer (TNBC) patients is hindered by a high incidence of chemoresistance (∼50%) (1, 2). The prevailing theory is that resistance and subsequent recurrence is driven by cancer stem cells. Unfortunately, the characterization of cancer stem cells is still largely based on correlative expression of cell surface markers, leaving the understanding of the molecular mechanisms incomplete. Several pathways are known to be intricately involved in the maintenance of both the stem/progenitor phenotype and tumorigenesis. Among these pathways, Wnt, FGF and Notch are some of most intensely studied (3–7). Additionally, p53 inactivation can lead to the generation of stem cell like signatures in tumor cells (8, 9).

Another confounding issue that has hampered the molecular characterization of TNBC cancer stem cells is the manner in which tumor initiation is studied and defined. The gold standard for tumor initiation is the ability to form a tumor in mice from low numbers (<1,000) of inoculating cells (10). However, the ability to form a detectable tumor is, at minimum, a two-step process. First, tumor-initiating cancer stem cells must form a nascent mass. The nascent mass must then expand and invade into the surrounding tissue to form a detectable mass. The expansion step is not mediated by cancer stem cell proliferation, but rather by partially differentiated transit amplifying cells, analogous to the process in developing tissue (11). Thus, it is also critical to delineate the molecular distinctions between bona fide cancer stem cells and more differentiated transit amplifying cells.

Tumor initiating cells have been defined using both genetically engineered mouse models and by isolation and characterization of human epithelial cells (12, 13). In one case, cells capable of forming K-Ras driven tumors in the lungs of mice were identified by ablation using bleomycin (13). In the other, human mammary epithelial cells were isolated from normal woman using defined conditions, which allowed the cells to maintain both luminal and basal characteristics and cell surface markers (12). These findings indicate that only a subset of cells in a given tissue, with progenitor-like potential, have the ability to give rise to tumors, while others, even with sufficient driver mutations, do not.

Thus, one of the necessary steps in functionally evaluating differences between tumor-initiating cells is defining the cell of origin for carcinomas. Presently, different methods are used to isolate cells from normal tissue that preferentially give rise to tumors (12, 13). These methods rely on identification/isolation of cells based on cell surface markers or outgrowth of primary cultures in defined media(12, 13). However, the ability to functionally define and prospectively isolate pure populations of tumor-initiating, and cancer stem, cells has not yet been achieved. In this report we report of the identification of a novel hierarchy of cancer stem cells. In this hierarchy, we demonstrate that differentiation status is based mechanistically on a non-canonical, cleavage-independent, Notch1 signaling pathway, resulting in the downregulation of the p53-deacetylase SIRT1, which is required for the fundamental cancer stem cell properties of tumor initiation and resistance to chemotherapy induced apoptosis.

## Results

### p53 is differentially acetylated in mammary progenitor cells and myoepithelial cells

We previously demonstrated that the p53 target gene thrombospondin-1 (Tsp-1) is differentially regulated in myopethelial-like human mammary epithelial cells (HMECs) compared to human fibroblasts (14). We sought to determine whether Tsp-1 was similarly regulated in bipotential progenitor-like mammary epithelial cells, (BPECs) (12). Notably, when BPECs are transformed via retroviral transduction of hTERT, SV40 Large T antigen (LT), and H-RasV12G they form tumors with 10,000-fold greater efficiency than commercially available human mammary epithelial cells (HMECs) transformed with the same set of genes (12).

To that end, we examined the expression of Tsp-1 in both HMECs and BPECs expressing LT alone and LT plus Ras. We found that ectopic expression of LT in tumor-initiating BPECs is sufficient to repress Tsp-1 (Figure 1A). Strikingly, the activity of LT on Tsp-1 expression in BPECs is similar to that observed in human fibroblasts and opposite to that of differentiated human epithelial cells(15).

**Figure 1.**
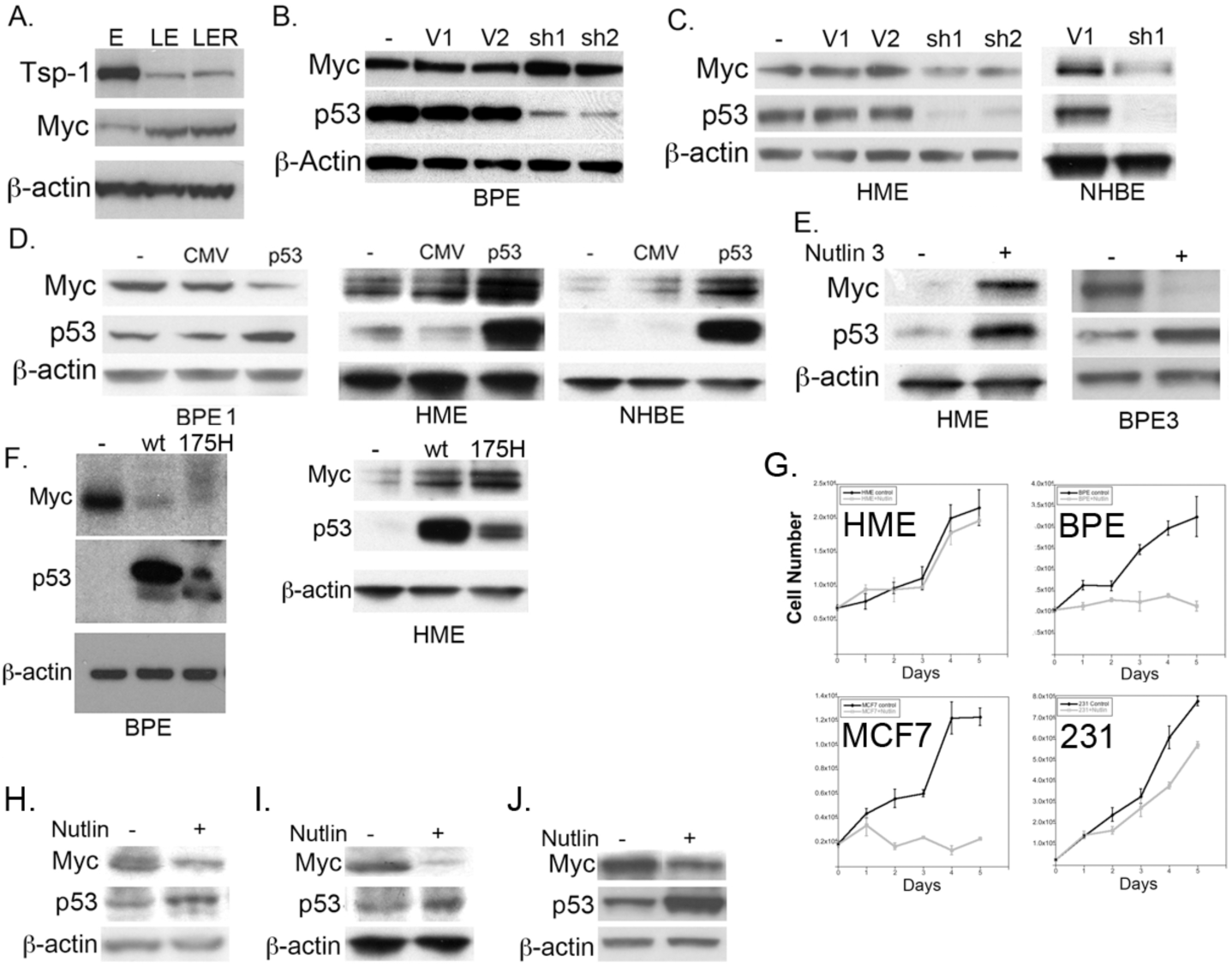
Differential effects of p53 activity in different cell types. Immunoblot analysis of: **(A)** Tsp-1, *c-myc* and actin expression in BPE cells in the presence and absence of ectopic expression of SV40 Large T antigen (LT) (BPLE), and HRasV12 (BPLER); **(B)** Myc, p53 and actin expression in wild type BPE cells transduced with vector controls pLKO.1-puro (V1), pMKO-neo (V2), pLKO.1shp53 (sh1) and pMKOneoshp53 (sh2); **(C)** Myc, p53 and actin expression in normal HME, normal human bronchial epithelial cells (NHBE) transduced with vector controls pLKO.1-puro (V1), pMKO-neo (V2), pLKO.1shp53 (sh1) and pMKOneoshp53 (sh2); **(D)** Myc, p53 and actin expression in BPE1, HME, and NHBE cells that were transiently transfected with pCMVneo or pCMVneo-p53 (p53); **(E)** Myc, p53 and actin expression in HME and BPE3 cells that were untreated (-) or treated with 25μM Nutlin-3: **(F)** Myc, p53 and actin expression in BPE and HME cells that were transiently transfected with pCMVneo, pCMVneo-p53 (wt) or pCMVneo-p53R175H; **(G)** Growth curve of HME, BPE, MCF-7 and MDA-MB-231 (231) cells that were untreated (black line) or treated with 25μM Nutlin-3/day for 5 days (Grey line); **(H)** Western blot of Myc, p53 and actin expression in A549 lung cancer cells that were untreated (-) or treated with 25μM Nutlin-3; **(I)** Western blot of Myc, p53 and actin expression in LNCaP prostate cancer cells that were untreated (-) or treated with 25μM Nutlin-3; **(J)** Western blot of Myc, p53 and actin expression in HCT116 colon cancer cells that were untreated (-) or treated with 25μM Nutlin-3.

Because p53 stimulates Tsp-1 production in fibroblasts(16) we sought to determine whether the effects of LT were due to the inactivation of p53. Accordingly, we examined the expression of two established p53 target genes, *cd44* and *c-myc* (17, 18), in both BPE and HME cells. We transduced human mammary epithelial cells (HMEC’s), normal human bronchial epithelial (NHBE) cells, BPECs and human dermal fibroblasts with lentiviral constructs specifying two independent shRNA sequences specific for p53: one that targets the coding sequence (sh1) (19), and a second that targets the 3’ UTR (sh2). We observed, via western blot analysis, that both shRNA sequences reduced p53 protein levels by greater than 90% (Figure 1B). We confirmed the loss of p53 function in these cells by examining the expression of the p53 target gene, p21(18, 20). Western blot analysis confirmed that p21 expression in these cells was greatly reduced and was not induced following treatment with the DNA damaging agent etoposide (Supplemental Figure S1). Consistent with the effects of LT expression, we observed that silencing of p53 had different effects on Myc expression in the different cell types. Silencing p53 in in BPECs resulted in increased Myc levels (Figure 1B). However, in HMECs and NHBECs silencing p53 resulted in ∼3-fold repression of *c-myc* (Figure 1C).

We then sought to determine whether these results could be confirmed by augmenting endogenous p53 activity without genotoxic stress or ectopic expression. Accordingly, we used the MDM2 (mouse double minute 2 homolog) inhibitor Nutlin-3, which increases p53 protein stability (21). Consistent with the effects of ectopically expressing p53, treatment of two independently isolated lines of BPECs with Nutlin-3 resulted in ∼7-fold decrease in Myc protein levels (Figure 1D), and treatment of human dermal and lung fibroblasts with Nutlin-3 resulted in a similar decrease in Myc protein levels, as determined by western blot analysis (Figure S2). Conversely, treatment of HMEC’s with Nutlin-3 resulted in a 8.5-fold *increase* in Myc protein levels (Figure 1E). We confirmed by both RT-PCR and ChiP (chromatin immunoprecipitation) that the effects of p53 on Myc expression were at the transcriptional level (Figure S3). Thus, regulation of Myc expression by p53 in tumor-initiating BPEC’s is more like mesenchymal cells than differentiated epithelial cells.

### Inhibition of p53 has different effects on cellular proliferation in different cells

We then examined whether these striking differences in p53 activity had any functional consequences on the two different types of epithelial cells. We tested the effects of Nutlin-3 on the proliferation of BPE and HME cells and the breast cancer cell lines, MCF-7 and MDA-MB-231. Nutlin-3 had little to no effect on the proliferation of HME cells but significantly inhibited the BPE cell proliferation (Figure 1F). Moreover, Nutlin-3 treatment significantly inhibited the growth of MCF-7 cells, which express wild-type p53 (Figure 1F). Conversely, Nutlin-3 treatment had little to no effect on the proliferation of MDA-MB-231 cells, which express a mutant p53 protein that is not a substrate for MDM2 (Figure 1F).

These results suggest that loss of p53 activity in tumor-initiating cells would result in increased proliferation and expansion of premalignant cells, while loss of p53 activity in differentiated epithelial cells would likely have little to no effect on their proliferative capacity.

### p53 is differentially modified in different cell types

Having demonstrated that modulation of p53 expression/activity had different effects in progenitor-like and myoepithelial-like mammary epithelial cells, we sought to ascertain whether this was the result of differential modification of the p53 protein itself in these cell types. Thus, we examined p53 post-translational modification in these cell types. We focused on two types of modifications that have been demonstrated to affect p53 transcriptional activity: phosphorylation and acetylation (22–24). We observed no differences in p53 phosphorylation at serines 15 and 20, nor did we observe differences in p21 expression, upon treatment of BPEC’s and HMEC’s with nutlin-3 (Figure S4). We then measured p53 acetylation levels, as residues 373 and 382 are acetylated by p300 and deacetylated by SIRT1 (23, 25). Contrary to the observations with regard to phosphorylation, Nutlin-3 treatment induced an ∼20-fold increase in the levels of acetylated p53 in BPECs but virtually no change in p53 acetylation in HMECs despite comparable increases in total p53 levels (Figure 2A).

**Figure 2.**
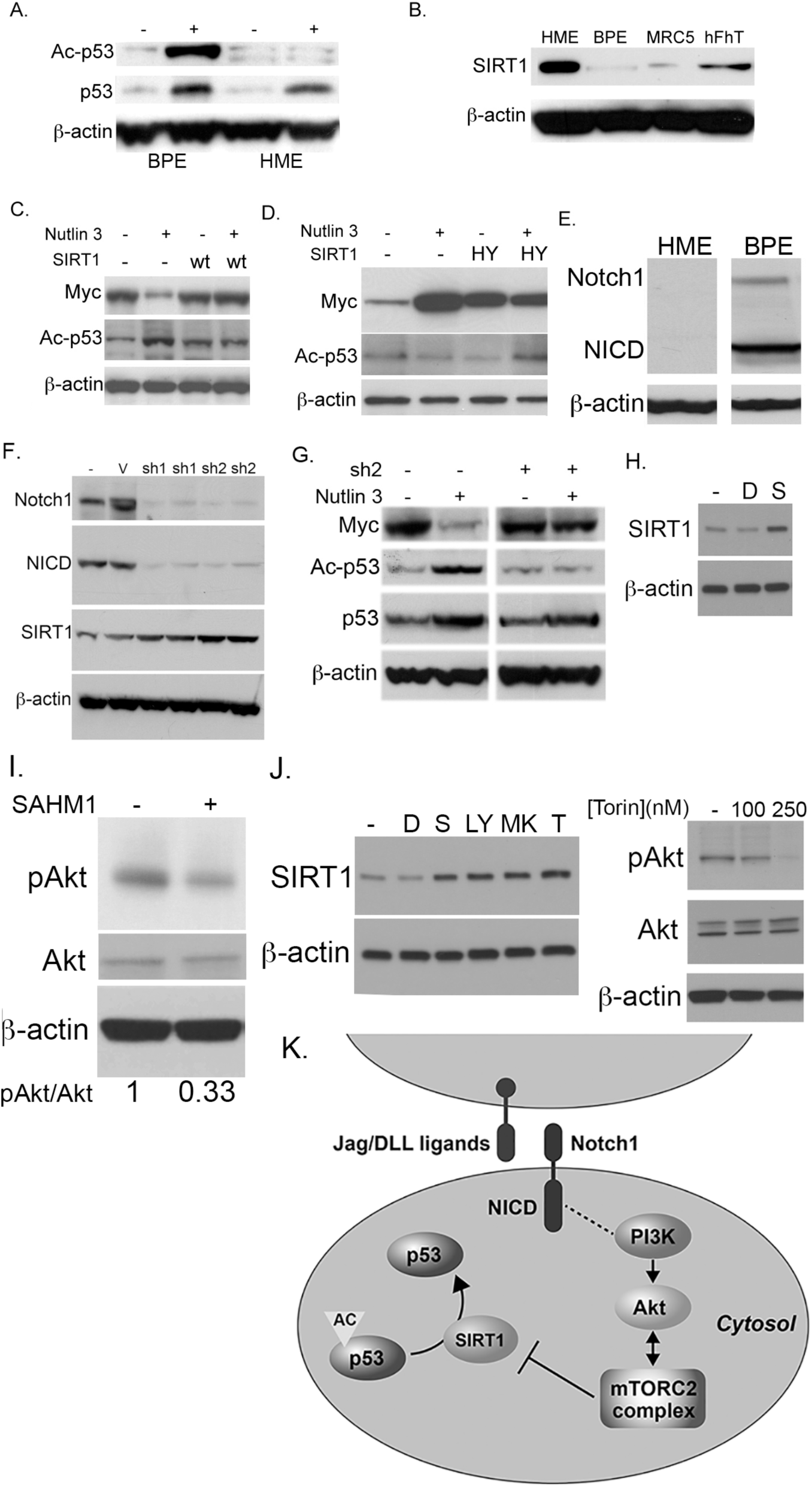
p53 acetylation is regulated by the non-canonical Notch1 pathway. Immunoblot analysis of: **(A)** Acetylated p53 (Ac-p53), total p53 and actin expression in BPE and HME cells that were untreated (-) or treated with Nutlin-3 for 8 hours; **(B)** SIRT1 and actin expression in HME, BPE, lung MRC5 fibroblasts and hFhT cells; **(C)** Myc, acetylated p53 (Ac-p53) and actin expression in BPE cells transduced with vector alone (-), or wtSIRT1 (wt) that were untreated (-) or treated with 25μM Nutlin-3 for 8 hours (+); **(D)** Myc, acetylated p53 (Ac-p53) and actin expression in HME cells transduced with vector alone (-) or SIRT1-HY (HY) that were untreated (-) or treated with 25μM Nutlin-3 for 8 hours (+); **(E)** Notch1, NICD and actin expression in HME and BPE cells; **(F)** Notch1, NICD, SIRT1 and actin expression in wild type BPE and cells transduced with two Notch1 shRNAs; **(G)** Myc, Ac-p53, p53 and actin expression in BPE and BPE-shNotch cells that were untreated (-) or treated with Nutlin-3 (+) for 8 hours; **(H)** SIRT1 and actin expression in BPE cells that were untreated (-) or treated with 25µM DAPT (D) and15μM SAHM1 (S) for 24 hours; **(I)** pAKT, total Akt, and actin expression in BPE cells that were untreated (-) or treated with 15μM SAHM1 (+) for 1 hour; **(J) (Left)** SIRT1 and actin expression in BPE cells that were untreated (-) or treated with 25μM DAPT (D), 15μM SAHM1 (S), 10μM LY294002 (LY), 10µM MK2206 (MK), or 250nM Torin (T); (**Right)** pAkt, total Akt and actin in BPE cells that were untreated (-) or treated with 100 or 250nM Torin; **(K)** Schematic diagram of Notch1 signaling pathway leading to the repression of SIRT1 expression.

To examine the cause of the different levels of p53 acetylation we measured the levels of SIRT1 expression in BPE and HME cells (25). By western blot analysis, we observed that BPECs express >10-fold lower levels of SIRT1 than HMECs (Figure 2B). In order to confirm that SIRT1 was mediating the different levels of p53 acetylation in these cells, we transduced BPECs and HMECs with retroviral constructs specifying wild-type SIRT1 and dominant-negative SIRT1 (SIRT1-HY) (Figure 2C, D) (25). We then treated cells with Nutlin-3 and analyzed p53 acetylation levels by western blot analysis. Treatment of BPEC-SIRT1 cells with Nutlin-3 completely inhibited p53 acetylation and concomitantly blocked the repression of Myc (Figure 2C). Conversely, treatment of HMECs expressing SIRT1-HY with Nutlin-3 resulted in increased levels of acetylated p53 and abrogated the stimulation of Myc (Figure 2D). These findings indicate that p53 acetylation, and transcriptional activity, is regulated differently in BPE and HME cells as a function of SIRT1 activity.

### Notch1 negatively regulates SIRT1 expression in tumor initiating and iPS cells

The Notch signaling pathway is a highly conserved pathway involved in multiple cellular and developmental processes. Significantly, the Notch pathway has been shown to be constitutively activated in different types of tumor initiating cells (26–28). The canonical Notch signaling pathway is activated by cell-cell interactions resulting in the cleavage of full-length, membrane-bound protein and subsequent release of Notch1 intracellular domain (NICD). The NICD translocates to the nucleus as part of a coactivator complex that targets CSL DNA-binding proteins. Alternatively, Notch1 can also signal through a non-canonical signaling pathway via PI3K/Akt/mTORC2 (29).

Based on reports that SIRT1 regulates the Notch1 pathway by either directly repressing Notch-activated genes or deacetylating NICD(30, 31), we examined whether Notch1 is involved in the SIRT1-mediated deacetylation of p53. We first measured Notch1 protein expression in both BPE and HME cells. We found, by western blot analysis, that BPE cells express dramatically higher levels of Notch1 than HME cells (Figure 2E).

To validate these findings and the role of Notch1 in SIRT1 expression, we transduced BPE cells with two lentiviral constructs specifying different Notch1 shRNA sequences (32). After validating that Notch1 levels were significantly reduced by both shRNA sequences (Figure 2F), we observed that silencing Notch1 resulted in the stimulation of SIRT1 protein levels (Figure 2F). We then treated the BPE-shNotch1 cells that displayed greater Notch1 knockdown (sh2) with Nutlin-3 and found that p53 acetylation levels were unchanged and Myc protein levels were no longer repressed (Figure 2G). These findings indicate that Notch1 is not only a target of SIRT1 but also represses its expression, implicating a feedback loop is present in these cells that regulates Notch1 and SIRT1 expression/activity.

To determine the mechanism of Notch1-mediated regulation of SIRT1 we treated BPE cells with both the Notch1/γ-secretase inhibitor DAPT and a stapled peptide mimetic of MAML1, SAHM1 (33). Treatment of BPE cells with DAPT resulted in reduced generation of NICD (Figure S6) but failed to inhibit the repression of SIRT1 (Figure 2H). Conversely, treatment of BPE cells with SAHM1 resulted in the derepression of SIRT1 (Figure 2H). This finding suggests that Notch1-mediated inhibition of the expression of SIRT1 is via the non-canonical signaling pathway(29) independent of NICD cleavage.

### Notch1 signals via PI3K/Akt/mTORC2 to repress SIRT1 expression

To test whether the non-canonical Notch pathway regulates SIRT1 expression we examined the phosphorylation status of Akt. Specifically, we treated cells with SAHM1 and examined the effect on Akt phosphorylation at S473. We found that inhibition of Notch1 via SAHM1 resulted in a decrease in Akt phosphorylation of 3-fold (Figure 2I). Additionally, treatment with inhibitors of Notch1 (SAHM1), PI3K (LY294002), and Akt (MK2206) resulted in the derepression of SIRT1 expression by 2.3-fold, 3.4-fold and 3.1-fold, respectively (P=0.038, 0.040, and 0.0095, respectively by Student’s t-test) (Figure 2J). We then assessed whether mTORC2 was involved in this pathway. To that end, we employed the mTORC1/2 dual inhibitor, Torin, as well as rapamycin as a control for inhibition of mTORC1 alone. Treatment of BPE cells with rapamycin had no effect on the expression of SIRT1 (Figure S7). Conversely, treatment with Torin resulted in a decrease in Akt phosphorylation of 3.3-fold (P=0.0149, by ANOVA), and a derepression of SIRT1 of 2.5-fold (P=0.009, by ANOVA) (Figure 3J right). These findings confirm that inhibition of Notch1 by SAHM1 inhibits the non-canonical Notch pathway, which derepresses SIRT1 expression via PI3K/Akt/mTORC2 signaling. Taken together, these results demonstrate that Notch1 negatively regulates SIRT1 expression in tumor initiating cells.

**Figure 3.**
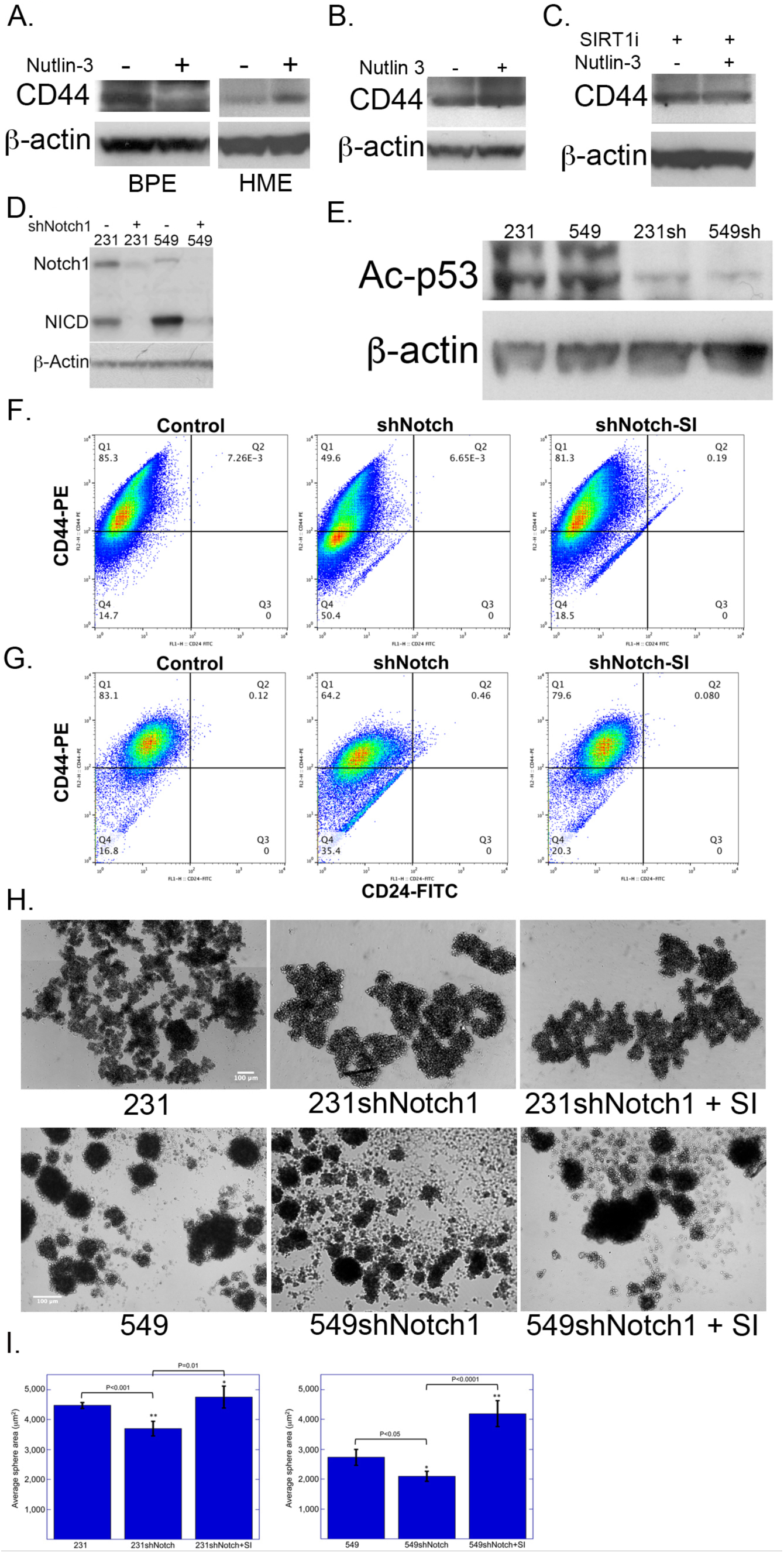
Effects of p53 acetylation on cancer stem cell properties *in vitro*. Immunoblot analysis of: **(A)** CD44 and β-actin expression in BPE and HME cells that were untreated or treated with 25μM Nutlin-3; **(B)** CD44 and β-actin expression in BPE-shNotch1 cells that were untreated or treated with 25μM Nutlin-3; **(C)** CD44 and β-actin expression in HME cells that were untreated or treated with 25μM Nutlin-3 in the presence of 20μM SIRT1 inhibitor for 7 hours; **(D)** Notch1, NICD and β-actin expression in wild-type MDA-MB-231 (231) and BT-549 (549) breast cancer cells and 231 and 549 cells in which Notch1 was silenced via shRNA; **(E)** Acetylated p53 and β-actin expression in wild-type MDA-MB-231 (231) and BT-549 (549) breast cancer cells and 231 and 549 cells in which Notch1 was silenced via shRNA; **(F)** FACS analysis of the CD44^hi^/CD24^lo^ populations in MDA-MB-231 breast cancer cells, in which Notch1 was silenced via shRNA (shNotch1), and treated with 20μM SIRT1 inhibitor (shNotch-SI) overnight prior to FACS sorting; **(G)** FACS analysis of the CD44^hi^/CD24^lo^ populations in BT-549 breast cancer cells, in which Notch1 was silenced via shRNA (shNotch1), and treated with 20μM SIRT1 inhibitor (shNotch-SI) overnight prior to FACS sorting; **(H)** Light field microscopy of tumorspheres formed by MDA-MB-231 (231) and BT-549 (549) breast cancer cells, 231 and 549 cells in which Notch1 was silenced via shRNA, and 231- and 549shNotch1 cells treated with 20μM SIRT1 inhibitor daily for 12 days; **(I)** Plot of average area (μm^2^) of tumorspheres formed by MDA-MB-231 (left) and BT-549 cells (right) depicted in Figure 4H.

Finally, we examined whether the regulation of SIRT1 expression by Notch1 signaling was at the transcriptional or post-transcriptional level. We performed reverse transcriptase (RT) PCR on BPE cells that were untreated or treated with SAHM1 and found that there was no change in RNA levels of SIRT1 (Figure S8). These findings led us to test whether Notch1 signaling affected the degradation of SIRT1 as SIRT1 protein levels have been shown to be regulated by proteasomal degradation (34). We treated BPE cells with the proteasome inhibitor MG132 and analyzed protein levels via western blot. We found that MG132 treatment induced a significant increase in SIRT1 protein levels (Figure S8). Taken together these results indicate the down-regulation of SIRT1 by the non-canonical Notch1 signaling pathway is mediated by proteasomal degradation (Figure 2K).

The findings reported here indicate that BPE cells resemble mesenchymal cells with regard to the activity of wild-type p53 and epithelial cells with regard to the activity of mutant p53. Moreover, BPE cells express markers consistent with mammary epithelial progenitor cells, while HME cells express markers consistent with myoepithelial cells (12). Based on these observations we speculated that the differential regulation of p53 is a function of the differentiation status of epithelial cells. To test the hypothesis we examined the expression of SIRT1, p53, and Notch1 in human inducible pluripotent stem cells (iPSc’s) that were derived by ectopic expression of Myc, Oct4, KLF4 and Sox2 in normal human keratinocytes (35). We found that human iPSc’s expressed very low levels of SIRT1 and high levels of Notch1, similar to tumor-initiating cells (Figure S9). Moreover, when these iPSc’s cells were treated with the Notch1 inhibitor SAHM1, we observed that SIRT1 levels were stimulated (Figure S9). Finally, when these cells were treated with Nutlin-3, p53 levels were increased and Myc levels were decreased, again, consistent with the results observed with mammary epithelial progenitor cells (Figure S9). These findings suggest that Notch1 and SIRT1 levels, as well as acetylation of p53, are conserved between progenitor/tumor initiating cells and pluripotent stem cells.

### Notch mediated acetylation of p53 regulates the cancer stem cell population

Having demonstrated that Notch modulates p53 activity in mammary epithelial progenitor-like cells and pluripotent iPS cells, we sought to determine whether it also functionally regulated the progenitor phenotype in mammary epithelial cells. We therefore turned our attention to the stem cell marker CD44, which is negatively regulated by p53 in BPE cells (17). To determine whether CD44 expression was regulated in an analogous manner in normal HME cells we treated both HME and BPE cells with Nutlin-3 and examined its expression by western blot analysis. Consistent with published reports, we observed that CD44 expression was repressed following Nutlin-3 treatment of BPE cells (Figure 3A). Conversely, in HME cells, Nutlin-3 treatment resulted in CD44 upregulation (Figure 3A). In order to determine whether the cell specific differential regulation of CD44 was mediated by Notch1 signaling we examined the effects of Nutlin-3 treatment in BPE-shNotch1 cells. We found that in the absence of Notch1, Nutlin-3 treatment no longer resulted in the repression of CD44, but rather a modest stimulation of CD44 expression (Figure 3B). We then treated HME cells with Nutlin-3 in the presence and absence of the SIRT1 inhibitor, 6-Chloro-2,3,4,9-tetrahydro-1H-carbazole-1-carboxamide (SIRT1i) (36). Western blot analysis revealed that inhibition of SIRT1 blocked the Nutlin-3 induced stimulation of CD44 in HME cells and led to the repression of CD44, similar to the effects in BPE cells (Figure 3C).

Based on these findings we sought to determine whether CD44 was regulated via the same mechanism in breast cancer cells harboring a p53 mutation. Accordingly, we examined two triple negative breast cancer cell lines with two different p53 mutations: MDA-MB-231 (R280K) and BT-549 (R249S) (37). We sought to determine whether these mutant p53 proteins were, similar to wild-type p53, also acetylated in a Notch1-dependent manner. Accordingly, we silenced Notch1 expression in these cell lines via lentiviral transduction of a vector specifying an shRNA sequence specific for Notch1. Western blot analysis confirmed that Notch1 was silenced by >95% in 231 and 549 cells (Figure 3D). Western blot analysis further revealed that p53 acetylation levels were decreased ∼5-fold in both 231 and 549 cells following silencing of Notch1 (Figure 3E), indicating that the Notch1-SIRT1-p53 pathway was active in breast cancer cells in addition to mammary epithelial cells.

To determine whether SIRT1 regulated CD44 expression in breast cancer cells, as observed in BPE cells, we FACS sorted these cells to determine the effect of Notch silencing on the cancer stem cell population, as defined by high expression of CD44 and low expression of CD24 (CD44^hi^/CD24^low^) (10). FACS analysis revealed that Notch1 silencing resulted in a reduction in the CD44^hi^ cancer stem cell population of both 231 and 549 cells (85.3% to 49.6% and 83.1% to 64.2%, respectively) (Figure 3F and G). The effects of silencing Notch1 were abrogated by treating cells with a chemical inhibitor of SIRT1 (Figure 3F and G).

Finally, to determine whether the change in cancer stem cell marker profiles correlated with a phenotypic change in functional cancer stem cells, we performed tumor sphere assays with the MDA-MB-231 and BT-549 cell lines (38). Consistent with the FACS analysis, we found that the number of tumor spheres formed by both 231 and 549 cells was dramatically decreased when Notch1 expression was silenced (Figure 3H and I). Strikingly, when SIRT1 activity was inhibited in Notch1 silenced cells the average size of the tumor spheres increased by 125% for 231 cells and 198% for 549 cells, while the number of spheres formed was not significantly altered (Figure 3H and I). These results suggest that Notch1 mediates two aspects of tumor sphere formation: size, which is SIRT1/p53 dependent; and number, which is SIRT1/p53 independent. These findings indicate that mutant p53 functions in the opposite manner as wild-type p53 physiologically as well as biochemically. Specifically, Notch1-mediated acetylation of wild-type p53 represses CD44 and the progenitor-like properties of mammary epithelial cells, whereas it stimulates CD44 and augments the cancer stem cell properties in breast cancer cells harboring mutant p53.

### The canonical breast cancer stem cell population is comprised of two distinct populations

Having demonstrated that Notch modulates p53 activity in mammary epithelial progenitor-like cells and pluripotent iPS cells, we sought to determine whether it also functionally regulated the progenitor phenotype in breast cancer stem cells. Accordingly, we examined whether Notch1 expression could be used, in conjunction with CD44, CD24 and ESA to isolate cancer stem cells. When we examined Notch1 expression via FACS in the CD44^hi^/CD24^low^/ESA^+^ population we made an unexpected observation. We found, that both cell lines contained two distinct populations of cells within the CD44^hi^/CD24^low^/ESA^+^ population that could be distinguished by the level of Notch1 expression (Figure 4A and S8). The main population of cells expressed very low levels of Notch1, while a side population of cells expressed ∼1,000-fold higher levels of Notch1 (Figure 4B and S10).

**Figure 4.**
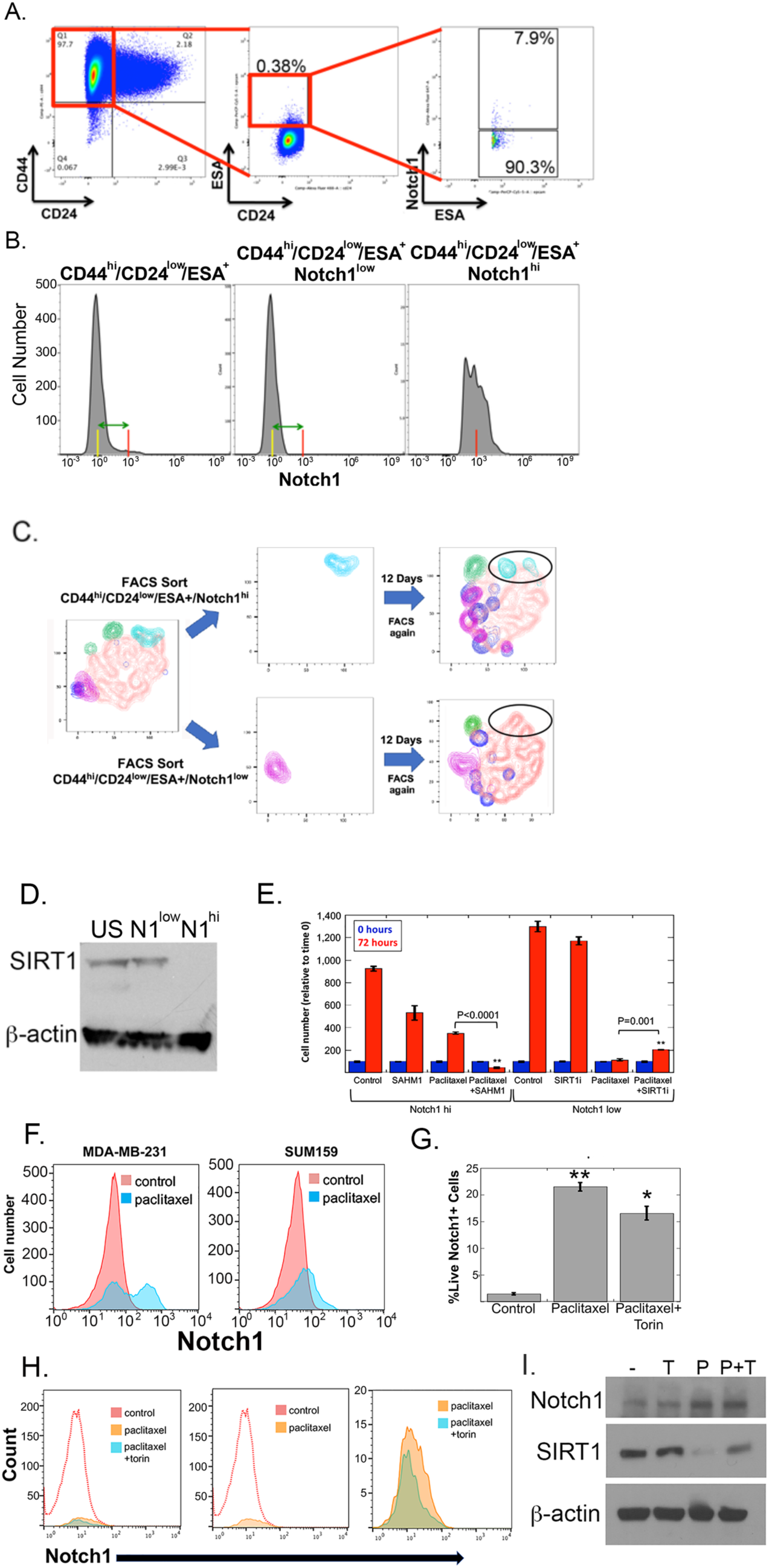
Role of Notch1 in tumor initiation by breast cancer stem cells. **(A)** FACS analysis of the Notch1 expression in CD44^hi^/CD24^lo^/ESA^+^ populations in SUM159 breast cancer cells; **(B)** Histogram of FACS analysis of Notch1 expression in CD44^hi^/CD24^lo^/ESA^+^, CD44^hi^/CD24^lo^/ESA^+^/Notch1^low^ and CD44^hi^/CD24^lo^/ESA^+^/Notch1^hi^ cells in SUM159 cells; **(C)** tSNE analysis of cell populations present before FACS sorting and after 12 days in culture **(D)** Immunoblot analysis of SIRT1 and β-actin expression in unsorted (US), CD44^hi^/CD24^lo^/ESA^+^/Notch1^low^ (N1^low^), and CD44^hi^/CD24^lo^/ESA^+^/Notch1^hi^ (N1^hi^) SUM159 cells; **(E)** Plot of cell number of CD44^hi^/CD24^lo^/ESA^+^/Notch1^high^ cells following treatment with SAHM1, paclitaxel, and SAHM1+paclitaxel, as well as and Notch1^low^ cells following treatment with SIRT1 inhibitor, paclitaxel and SIRT1i+paclitaxel; **(F)** Histogram of Notch1 expression vs number of live cells following treatment with saline (control) or paclitaxel; **(G)** Plot of percentage of live SUM159 Notch1+ cells following treatment with saline (control), 20nM paclitaxel for 48 hours, or 20nM paclitaxel for 48 hours + 250nM Torin for 16 hours; **(H)** Histogram of live SUM159 Notch1+ cells following treatment with saline (control), 20nM paclitaxel for 48 hours, or 20nM paclitaxel for 48 hours + 250nM Torin for 16 hours; **(I)** Western blot analysis of Notch1, SIRT1 and β-actin expression in SUM159 cells following treatment with saline (-), 250nM Torin (T), 20nM paclitaxel for 48 hours (P), or 20nM paclitaxel for 48 hours + 250nM Torin for 16 hours (P+T);

To determine if these two populations of cells had more characteristics associated with stem cells we performed a differentiation experiment by growing each population of cells in culture for 13 days following FACS sorting. After thirteen days in culture we analyzed the composition of each population of cells by flow cytometry by tSNE analysis. We found that the Notch1^hi^ cells gave rise to all of the population of cells that were present in the parental population before sorting with similar percentages (Figure 4C). Conversely, the Notch1^low^ cells were able to give rise to all of the populations present in the parent population with the exception of the Notch1^hi^ population (Figure 4C). These findings suggest that the Notch1^hi^ cells are more stem-like than the Notch1^low^ cells and sit above them in the hierarchy of differentiation.

Based on the findings in BPE cells and the tSNE analysis, we speculated that the Notch1^hi^ cells would express lower levels of SIRT1 than the Notch1^low^ cells. To test this, we measured the levels of SIRT1 by western blot in both populations, as well as in the unsorted population of SUM159 cells. We found that the Notch1^low^ cells expressed high levels of SIRT1, virtually indistinguishable from the unsorted population (Figure 4D). Conversely, the Notch1^hi^ cells expressed virtually undetectable levels of SIRT1 (Figure 4D).

A prominent and well-studied characteristic of cancer stem cells is an increased resistance to chemotherapy (39–41). With that in mind we sought to determine whether the Notch1^hi^ cancer stem cells were more resistant to chemotherapy than their Notch1^low^ counterparts. Specifically, we examined the response to paclitaxel, as it represents one of the most commonly utilized therapeutic agents for TNBC, taxanes. Accordingly, we treated the high Notch1 expressing cells with the Notch inhibitor, SAHM1, alone and in combination with paclitaxel. Analogously, we treated the low Notch1 cells with the SIRT1 inhibitor, EX527, alone and in combination with paclitaxel. We found, in the absence of treatment, that after 72 hours the low Notch1 cells had proliferated significantly faster than the high Notch1 cells (13 vs 9-fold, P=0.003 by ANOVA) (Figure 4E). We observed that treatment with the Notch1 inhibitor SAHM1 reduced the proliferation rate of the Notch1^hi^ cells by 40% (9-fold vs 5.3-fold increase in cell number; P=0.005 by ANOVA) (Figure 4E). Conversely, inhibition of SIRT1 activity did not significantly affect the proliferation of the Notch1^low^ cells (Figure 4E).

However, treatment with 20nM paclitaxel had strikingly different effects on the two cell types. Specifically, the Notch1^hi^ cells continued to proliferate (3.5-fold; P<0.0001 by ANOVA) in the presence of paclitaxel, while the proliferation of Notch^low^ cells was significantly inhibited after the 72 hour treatment period (11.4-fold; P<0.0001 by ANOVA) (Figure 4E). Strikingly, when the Notch1^hi^ cells were treated with paclitaxel in the presence of SAHM1, the cell number decreased by 55%, indicating that inhibiting Notch1 activity rendered these cells sensitive to killing by paclitaxel (Figure 4E). Conversely, when the low Notch1 expressing cells were treated with paclitaxel in the presence of EX527, the cell number increased by 2-fold, indicating that inhibiting SIRT1 rendered these cells resistant to paclitaxel-mediated cell killing (Figure 4E).

We then treated an unsorted population of SUM159 and MDA-MB-231 cells with paclitaxel to examine the effects of treating a heterogeneous tumor. Following treatment with paclitaxel for 72 hours we examined the percentage of low and high Notch1 expressing cells by FACS. We found that the percentage of high Notch1 cells dramatically increased in both cell types following treatment with paclitaxel (MDA-MB-231 4.1 vs 52.6; P<0.001 by ANOVA; SUM159 2.1 vs 20.5 P<0.005 by ANOVA) (Figure 4F). These results indicate that high Notch1 expressing cells possess an intrinsic chemoresistance characteristic of cancer stem cells.

To follow up on these findings we sought to determine whether inhibiting mTORC2 activity would have similar results on the chemoresistance of high Notch1 expressing breast cancer cells. To that end we treated SUM159 cells with paclitaxel, as described above, for 48 hours, the last 16 of which were in the presence of the mTORC1/2 inhibitor, Torin. FACS analysis revealed that the percentage of live high Notch1 expressing cells increased from ∼1.5% to ∼20% following paclitaxel treatment (P<0.001 by ANOVA) (Figure 4G and H). Consistent with the ability of Torin to inhibit the repression of SIRT1, treatment with Paclitaxel plus Torin resulted in an ∼20% reduction of live Notch1 high expressing cells (P=0.0167 by Student’s T-test).

Western blot confirmed that the levels of full-length Notch1 protein in the paclitaxel treated population of cells increased an average of 2.8-fold with a concomitant decrease of 10-fold in SIRT1 levels (Figure 4I). Moreover, while treatment of the parental population of SUM159 cells with Torin had no effect on the expression of SIRT1, treatment with Paclitaxel plus Torin resulted in a 4.1-fold (P=0.014 by ANOVA) increase in SIRT1 levels compared to paclitaxel alone (Figure 4I). These findings indicate that the chemoresistance of Notch1^hi^ expressing cells is mediated by the repression of SIRT1 via the Notch1/mTORC2 signaling pathway.

### Notch1 activity is mediated by p53 family members in breast cancer stem cells

We have demonstrated in mammary epithelial progenitor cells non-canonical Notch1 signaling mediates p53 activity. However, the vast majority of triple negative breast cancer cells, including those we have examined (MDA-MB-231, BT-549, and SUM159) all harbor mutations in the p53 gene. Interestingly, while MDA-MB-231 and BT-549 harbor gain of function mutations (R280K and R249S), SUM159 contains an insertion mutation of a leucine residue between V157 and R158, which is considered to be an inactivating mutation (42). Since Notch1-mediated inhibition of SIRT1 is required for chemoresistance (Figure 4E) we hypothesized that the functional target of SIRT1 may be a p53 family member rather than p53 itself.

Accordingly, we first examined the expression status of p63 and p73 in Notch1^hi^ and Notch1^low^ SUM159 cells. We found that expression levels of both p63 and p73, specifically the TA isoforms, were higher in Notch1^hi^ cells than in Notch1^low^ cells (Fig 5A). We followed up this observation by measuring p63 and p73 mRNA levels via RT-PCR following modulation of the Notch1 non-canonical pathway. Specifically, we measured the expression of the TA and DN isoforms of p63 and p73. These isoforms, differ in the N-terminal transactivation domain, which is present in the TA but absent in the DN isoforms, due to alternative promoter start sites (43). We found that in SUM159 cells, the TA isoforms of both p63 and 73 were decreased following inhibition of the non-canonical Notch1 pathway by IMR-1 in Notch1 high-expressing cells (Figure 5B). Consistent with these results, mRNA levels for these the TA isoforms were increased following inhibition of SIRT1 activity in the Notch1 low-expressing cells (Fig 5B). Conversely, expression of the DN isoforms were not affected by inhibition of SIRT1(Fig 5B). We also examined the regulation of p63 and p73 mRNA by Notch1 non-canonical pathway in MDA-MB-231 cells, and found that consistent with SUM159 cells, TAp73 is regulated by the Notch1—SIRT1 pathway, however, TAp63 expression was not affected (Supplemental 12). The difference in TAp63 regulation between these two cell lines may be due to the different p53 mutations in the two cell lines or an indication that the role of p63 in cancer stem cell function is not as significant as p73. Nevertheless, these results indicate that activation of the non-canonical Notch1 pathway leading to the degradation of SIRT1 increased transcription of TAp73 in two different TNBC cell lines.

**Figure 5.**
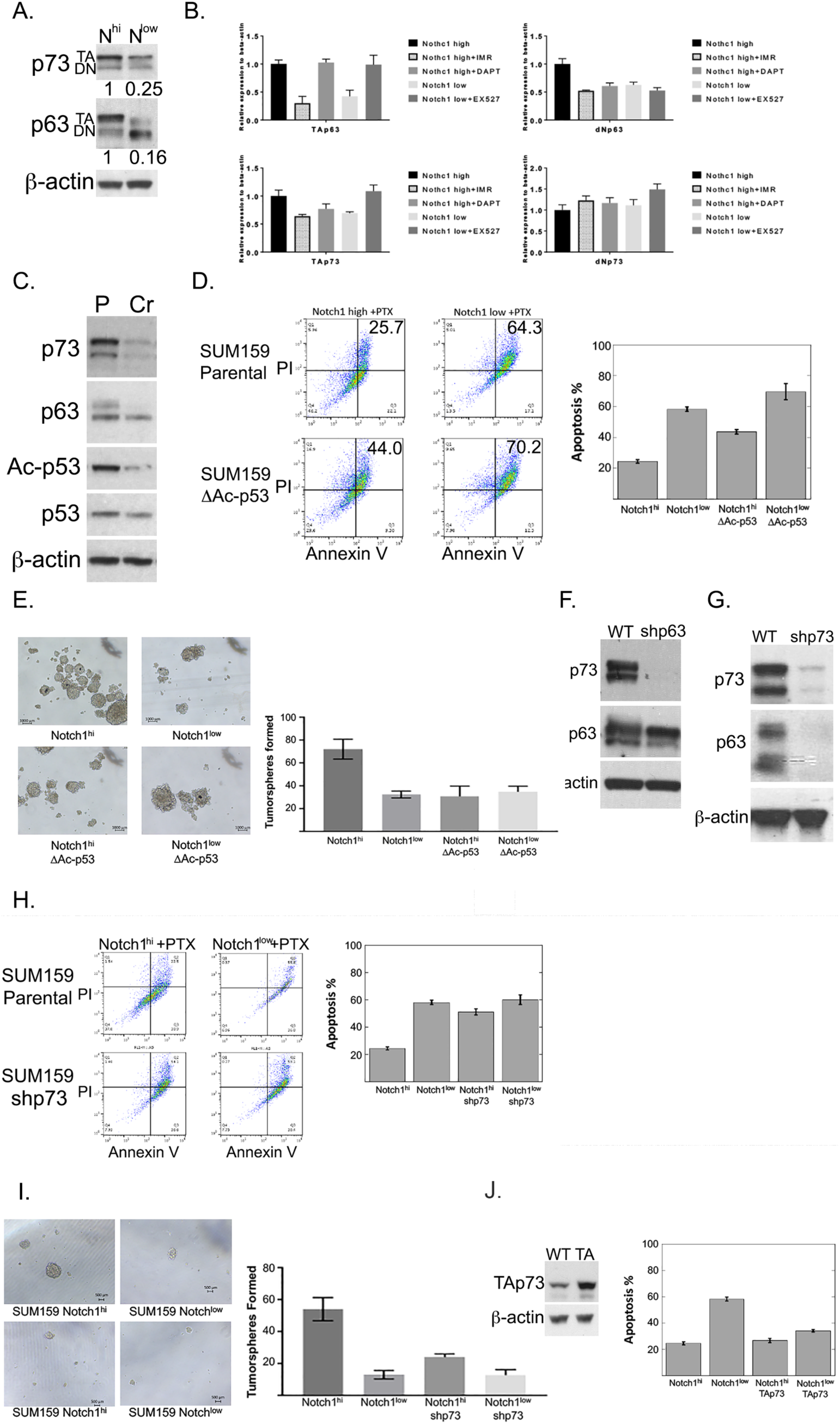
Role of p53 family members in cancer stem cell function. **(A)** Western blot of p73, p63, and β-actin in Notch1^hi^ and Notch1^low^ in SUM159 cancer stem cells; **(B)** RT-PCR of mRNA levels of TA and DN isoforms of p63 and p73 in Notch1^hi^ cells that were untreated or treated with SAHM1 and DAPT and Notch1^low^ cells that were untreated or treated with EX-527; **(C)** Western blot of p73, p63, acetylated p53 (Ac-p53), total p53 and β-actin in wild-type parental (P) SUM159 Notch1^hi^ cells and Notch1^hi^ cells in which the last 11 amino acids of the p53 protein had been deleted by CRISPR-Cas9 editing (Cr); **(D)** FACS analysis and bar graph of quantitation of apoptosis induced in SUM159 parental and CRISPR-edited, ΔAc-p53 cells, Notch1^hi^ and Notch1^low^ cells treated with 20nM paclitaxel; **(E)** Light-field images and bar graph of quantitation of tumorspheres formed by SUM159 parental and CRISPR-edited, ΔAc-p53 cells, Notch1^hi^ and Notch1^low^ cells; **(F)** Western blot of p73, p63, and β-actin in WT and shp63 SUM159 cells; **(G)** Western blot of p73, p63, and β-actin in WT and shp73 SUM159 cells; **(H)** FACS analysis and bar graph of quantitation of apoptosis induced in wild-type (parental) and shp73 SUM159 and cells, Notch1^hi^ and Notch1^low^ cells treated with 20nM paclitaxel; **(I)** Light-field images and bar graph of quantitation of tumorspheres formed by SUM159 parental and shp73, Notch1^hi^ and Notch1^low^ cells; **(J)** Western blot of TAp73 and β-actin in WT SUM159 and SUM159-TAp73 cells.

To determine whether p53 acetylation downstream from Notch1 signaling is required for the expression of TAp73, we generated cells lines in which p53 was unable to be acetylated at residue K382, using CRISPR-Cas9 editing. Specifically we utilized a gRNA that mediated a site specific double strand break at residue 379 resulting in a truncated form of p53 lacking the last 11 amino acids. The loss of acetylation at residue 382 was confirmed by western blot using an antibody specific for p53 acetylated at K328 (Fig 5C). We then measured the expression of both TAp73 and TAp63 in the CRISPR-edited cell lines and found that both mRNA and protein levels were significantly decreased compared to the parental cell line (Fig 5C).

To understand the functional role of ac-p53 (K382), we assessed the effects of acetylation on chemosensitivity/resistance of Notch1^hi^ and Notch1^low^ populations in parental SUM159 cells and SUM159 ac-p53 (K382)-null cells. We found that following treatment with paclitaxel apoptosis increased from 24.4% to 43.7% (P<0.0001 by ANOVA) in the Notch1^hi^ cells lacking ac-p53 (K382), compared to the parental SUM159 Notch1^hi^ cells (Fig 5D). We also examined the role of p53 acetylation in tumorsphere formation. Consistent with the chemoresistance analysis, we found that the number of tumorspheres formed by the Notch1^hi^ cells was decreased more than 2-fold when acetylation of p53 at residue 382 was disrupted by CRISPR-mediated gene editing (Fig 5E). Taken together the results of the chemoresistance and tumorsphere assays suggest that the indel mutation in p53 in SUM159 cells is not a complete loss of function mutation, as acetylation is required for Notch1-mediated chemoresistance and tumorsphere formation.

We then sought to determine whether p63 and p73 expression, activated by acetylation of p53, are functionally involved in Notch1-mediated cancer stem cell properties. Accordingly, we generated p63 and p73 knockdown cell lines via lentiviral transduction of shRNA. Knockdown of p63 and p73 was confirmed by western blot (Fig 5F and G). Surprisingly, we found that silencing of p73 also resulted in the downregulation of p63 (Figure 5F and G). These findings indicate the acetylated p53 stimulates the transcription of p73, which in turn stimulates the expression of p63.

Based on these findings, we performed chemoresistance and tumorsphere assays of parental SUM159 cells and SUM159 shp73. Consistent with SUM159 ac-p53 (K382)-null cells, we found that the Notch1^hi^ cells lacking p73 had increased sensitivity (24.4% to 51.1%, P<0.0001) to paclitaxel (Figure 5H). Moreover, knockdown of p73 resulted in ∼2-fold decrease in the number of tumorspheres formed by Notch1^hi^ cells compared to Notch1^hi^ parental cells (Figure 5I).

Finally, to determine whether TAp73 and TAp63 were necessary and sufficient for Notch1-mediated cancer stem cell function in SUM159 cells, we performed rescue experiments, in which we ectopically expressed TAp73 in SUM159 cells and then FACS sorted for Notch1^hi^ and Notch1^low^ cells. Following sorting we treated each population with paclitaxel (20nM) for three days and measured apoptosis by FACS staining of Annexin V and PI. We found that ectopic expression of TAp73 in Notch1^hi^ cells had no affect on their sensitivity to paclitaxel (Figure 5J). Conversely, ectopic expression of TAp73 in Notch1^low^ cells significantly decreased their sensitivity to paclitaxel (60.1% to 34.0%, P<0.0001, by ANOVA). Thus, our findings indicate that non-canonical Notch1-mediated degradation of SIRT1 increases acetylation of p53, which results in increased expression of TAp73, resulting in increased resistance to chemotherapy induced apoptosis.

### Notch1 repression of SIRT1 is required for tumor initiation

The results of the FACS sorting, chemoresistance and EMT transcription factor expression analyses raise the interesting hypothesis that only one of the two populations (Notch1^hi^ or Notch1^low^) within the CD44^hi^/CD24^low^/ESA^+^ “cancer stem cell” population represents functional, bona fide, cancer stem cells. To test this hypothesis, we collected the CD44^hi^/CD24^low^/ESA^+^/Notch1^hi^ and the CD44^hi^/CD24^low^/ESA^+^/Notch1^low^ cells following FACS sorting of SUM159 cells. Prior to sorting, the SUM159 cells were transduced with a lentiviral vector specifying firefly luciferase, allowing us to visualize tumor formation in real time. We allowed the cells to recover from the stress of sorting for 48 hours before injecting them into mice. We also performed a proliferation assay on the Notch1^hi^ and Notch1^low^ cells. Consistent with the results presented in Figure 5D, after 72 hours, the Notch1^low^ cells increased by ∼1300% relative to the starting number. Conversely the Notch1^hi^ cells increased in number by only ∼600% of the starting number, less than half the proliferation rate of the Notch1^low^ cells (Figure 6A).

**Figure 6.**
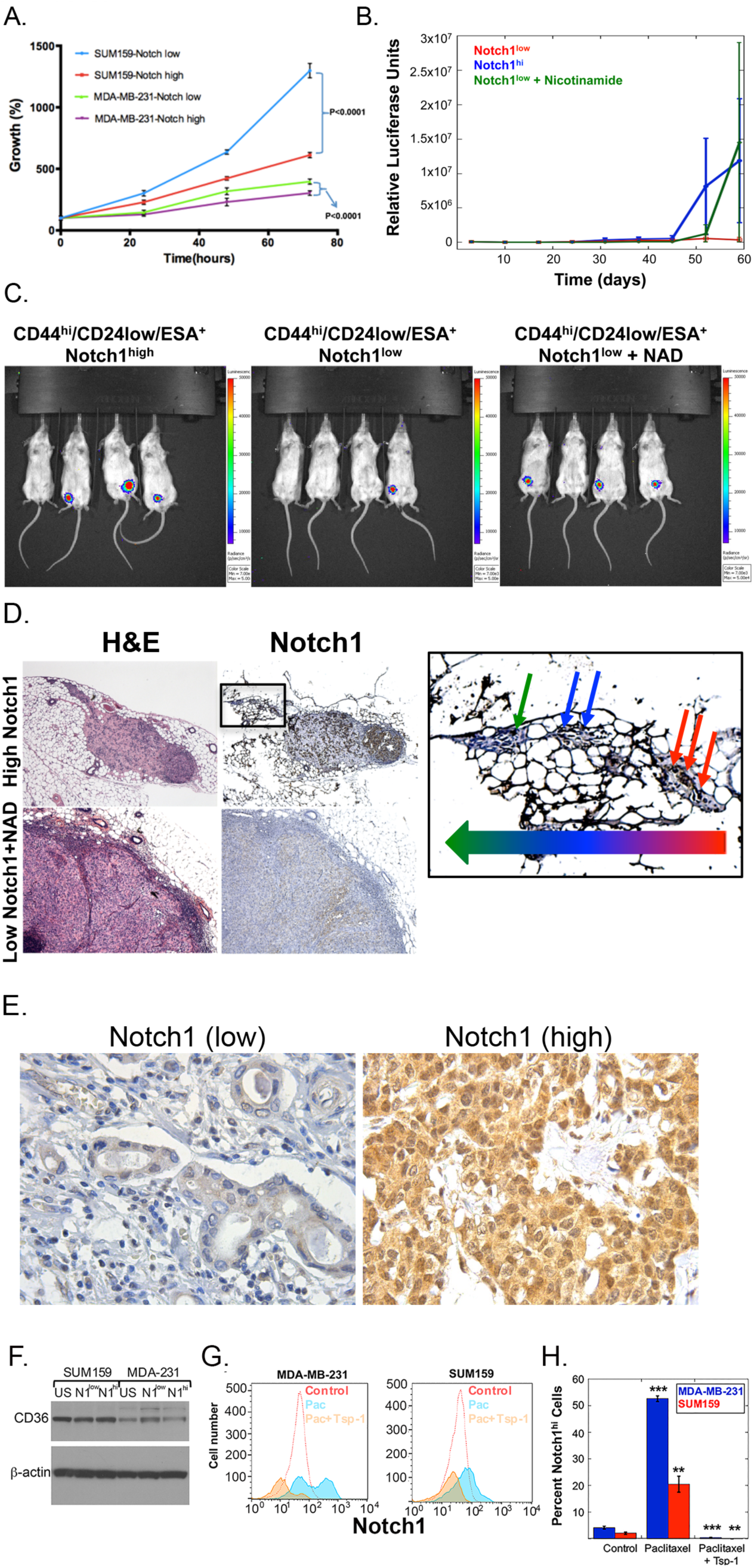
**(A)** Plot of cell number at day 0 and 48 hours after FACS sorting of CD44^hi^/CD24^lo^/ESA^+^/Notch1^hi^ and CD44^hi^/CD24^lo^/ESA^+^/Notch1^low^ cells from SUM159 vector control cells: **(B)** Plot of luciferase intensity of tumors formed by CD44^hi^/CD24^lo^/ESA^+^/Notch1^hi^ and CD44^hi^/CD24^lo^/ESA^+^/Notch1^low^ cells; **(C)** Images from Xenogen IVIS of mice injected with CD44^hi^/CD24^lo^/ESA^+^/Notch1^hi^ and CD44^hi^/CD24^lo^/ESA^+^/Notch1^low^ cells; **(D)** H&E and Notch1 staining (50X) of tumors formed by CD44^hi^/CD24^lo^/ESA^+^/Notch1^hi^ and CD44^hi^/CD24^lo^/ESA^+^/Notch1^low^ cells, boxed panel (left) is 4x magnification of boxed area in Notch1 IHC (small arrows depict Notch1 positive cells, large arrow depicts direction of tumor cell migration away from primary mass) (scale bars=100μm); **(E)** Notch1 staining of tumors from breast cancer patients. **(F)** Western blot analysis of CD36 and β-actin expression in unsorted (US), CD44^hi^/CD24^lo^/ESA^+^/Notch1^low^ (N1^low^), and CD44^hi^/CD24^lo^/ESA^+^/Notch1^hi^ (N1^hi^) MDA-MB-231 and SUM159 cells; **(G)** Histogram of Notch1 expression vs number of live MDA-MB-231 and SUM159 cells following treatment with saline (red outline), paclitaxel (blue) or paclitaxel followed by rhTsp1 (orange); **(H)** Graphical depiction of the percent live MDA-MB-231 and SUM159 Notch1^hi^ cells following treatment with saline (control), paclitaxel or paclitaxel followed by rhTsp1.

Following the 48-hour recovery time, we mixed the Notch1^hi^ and Notch1^low^ cells with Matrigel and injected 500 cells of each type into the mammary fat pad of SCID mice (n=8 for Notch1^hi^ and n=16 for Notch1^low^) (40). The mice injected with Notch1^low^ cells were divided into two cohorts (n=8/group). The first cohort was treated every day with saline, while the second was treated with the SIRT1 inhibitor nicotinamide (NAD) at a dose of 150mg/kg/day (44). Beginning on day 10 after inoculation, we monitored the growth of primary breast tumors using the Xenogen IVIS imaging system. After 60 days, 7/8 mice injected with the CD44^hi^/CD24^low^/ESA^+^/Notch1^hi^ cells developed detectable tumors, compared with only 2/8 mice injected with CD44^hi^/CD24^low^/ESA^+^/ Notch1^low^ cells (Figure 6B and C). Strikingly, 7/8 mice injected with CD44^hi^/CD24^low^/ESA^+^/Notch1^low^ cells that were injected with NAD also developed tumors.

Upon histological examination, we found that the Notch1^low^/NAD-treated tumors were comprised of cells expressing low or undetectable levels of Notch1 (Figure 6D), indicating that inhibition of SIRT1, functionally obviated the need for high levels of Notch1 expression for tumor initiation. Conversely, the tumors formed by the Notch1^hi^ cells contained a dense cluster of cells expressing high levels of Notch1 surrounded by cells expressing lower or undetectable levels of Notch1 (Figure 6D). Notably, the tumor cells that migrated away from the main tumor mass expressed lower levels of Notch1 (Figure 6D). These findings suggest that high levels of Notch1, or suppression of SIRT1 activity, are required for tumor initiation, but tumor expansion and invasion do not require high Notch1 levels. Moreover, these findings suggest that the Notch1^hi^ cells are functional cancer stem cells as they are able to differentiate into Notch1^low^ cells. Conversely, the Notch1^low^ cells treated with NAD gave rise to tumors comprised of cells expressing low to undetectable levels of Notch1. Taken together these findings demonstrate that CD44^hi^/CD24^low^/ESA^+^/ Notch1^hi^ cells are *bona fide* breast cancer stem cells and that the stem cell promoting properties of Notch1 are mediated by its repression of SIRT1.

### Notch1 expression and breast cancer patient progression

To determine whether the role of Notch1 in breast cancer development had any clinical relevance we examined a patient tumor tissue microarray (TMA) comprised of breast cancer tissue from 534 patients (Figure 6E). The TMA was stained for Notch1 expression and evaluated using the semi-quantitated Staining Index (SI) method (45). We found that Notch1 staining was significantly associated with larger tumors and high-grade cases (histologic grade III versus grade I-II, p=0.045) (Notch1 expression: low-grade cases, Notch1 SI=0-3, high-grade cases, SI=4-9) (Figure 6E and Table 1). These findings suggest that tumors with a greater percentage of high Notch1 expressing cells are more aggressive and imply that the number of cancer stem cells in a tumor correlates with the aggressiveness and progression of breast cancer.

**Table 1:**
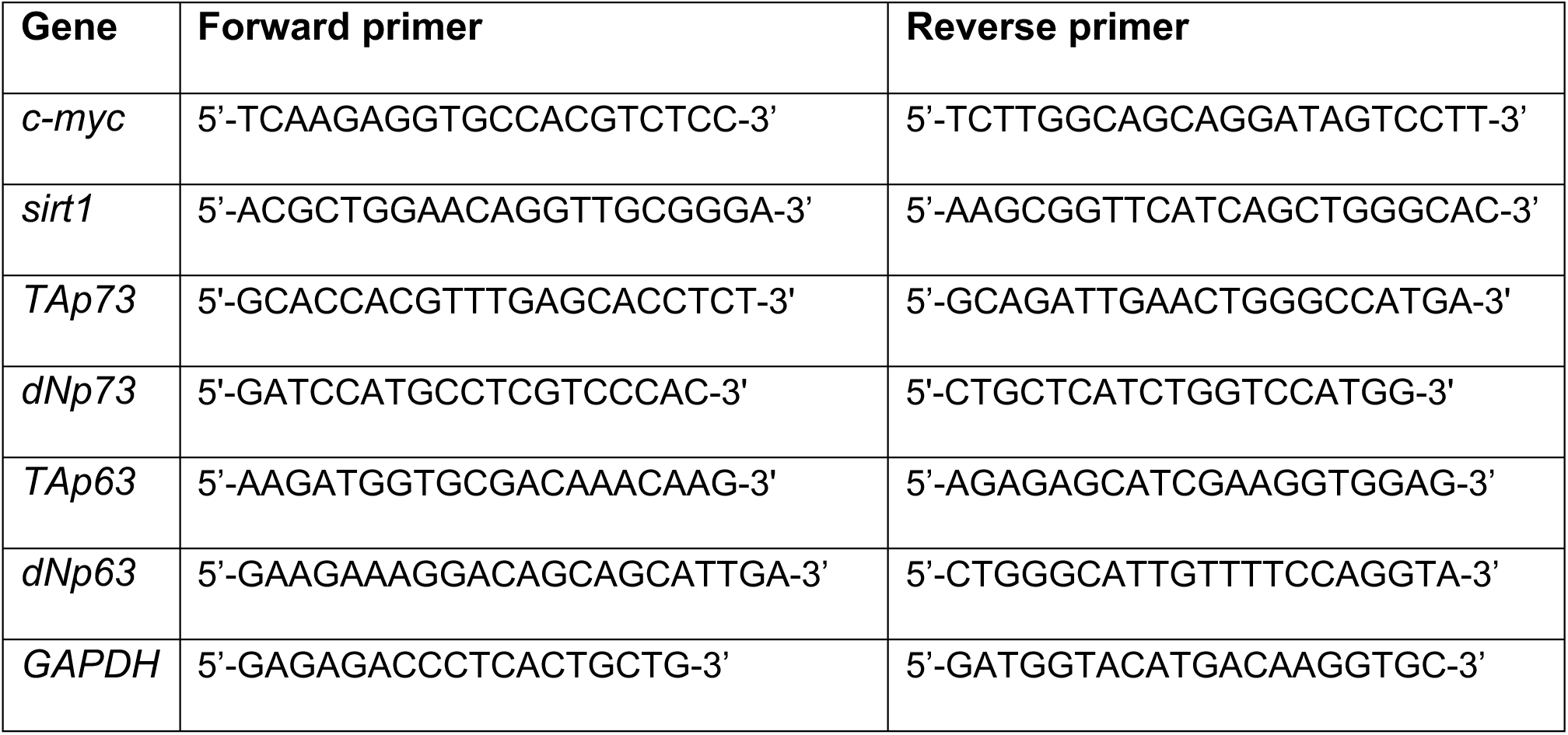
Sequence of the primers used for qPCR.

**Table 1.** Analysis of Notch1 expression in human breast cancer patients.

|  | Low Notch1 (SI 0-3) | High Notch1 (SI 4-9) | P-value |
| --- | --- | --- | --- |
| **Histologic Grade** |  |  |  |
| Grade 1-2 | 102/382 (27%) | 280/382 (73%) |  |
| Grade 3 | 11/71 (15.5%) | 60/71 (84.5%) | 0.045 |
| **Tumor Diameter** |  |  |  |
| <2cm | 20/44 (46%) | 24/44 (54%) |  |
| >2cm | 3/18 (16.7%) | 15/18 (83.3%) | 0.033 |

### Thrombospondin-1 specifically targets CD36 expressing cancer stem cells

The robust resistance to chemotherapy and potent tumor initiation potential, spurring the growth of recurrent tumors, of cancer stem cells adds an addition layer of complexity to the treatment of triple negative breast cancer patients. We have developed a therapeutic peptide derived from prosaposin, whose activity is mediated by the stimulation of thrombospondin-1 (Tsp-1) expression in MDSCs in the tumor microenvironment. The increased expression of Tsp-1 results tumor regression via the induction of apoptosis in ovarian cancer cells expressing CD36 (46). Thus, we sought to determine whether this therapeutic strategy could have applications for breast cancer. Therefore, we examined the expression of CD36 in unsorted SUM159 cells as well as the Notch1^low^ and Notch1^high^ populations. We found that CD36 was expressed by all populations, independent of Notch1 expression (Figure 6F). We then treated SUM159 and MDA-MB-231 cells that had been untreated or treated with paclitaxel for 72 hours with rhTsp-1 for 48 hours. We found that after treatment with paclitaxel the percentage of Notch1^hi^ cells increased from less than 3% in both MDA-MB-231 and SUM159 cells to over 50% in MDA-MB-231 cells and over 20% in SUM159 cells (Figure 6G and H). Strikingly, treatment with rhTsp-1 significantly appeared to specifically reduce the number, and percentage, of Notch1^high^ cells following paclitaxel treatment (Figure 6G and H). This finding indicates that breast cancer stem cells are sensitive to Tsp-1 and that a therapeutic agent that stimulates Tsp-1 would be effective at treating breast cancer alone, or in combination with paclitaxel.

## Discussion

We report here that breast cancer stem cell function is mediated by Notch1 activity via its repression of SIRT1 expression. Moreover, by sorting breast cancer cells for Notch1 expression, we were able to identify two populations of cells within the canonical stem cell population, defined as CD44^hi^, CD24^low^ and ESA^+^ (10). These populations differed in their level of Notch1 expression by 1,000-fold, based on FACS analysis. Strikingly, when injected into mice, 500 Notch1^hi^ cells were able to efficiently form tumors, while the same number of Notch1^low^ cells formed tumors at a significantly lower frequency. Consistent with the newly identified role of Notch1 in inhibiting SIRT1, treating a second cohort of mice injected with the Notch1^low^ cells with the SIRT1 inhibitor nicotinamide (thereby mimicking Notch1 function), resulted in a tumor initiation efficiency that was equivalent to the Notch1^hi^ cells. We also demonstrate that the repression of SIRT1 by Notch1 is mediated by the non-canonical pathway leading through PI3K/Akt/mTORC2 downstream from Notch1.

In keeping with findings that cancer stem cells have enhanced chemoresistant properties, we found that the Notch1/SIRT1 pathway is responsible for conferring resistance to traditional chemotherapy. Strikingly, despite being resistant to paclitaxel-mediated cell killing, we observed that the Notch1^hi^ are exquisitely sensitive to killing by thrombospondin-1 as they express the cell surface receptor, CD36, which mediates its pro-apoptotic activity. These findings represent the first demonstration of a distinct and necessary biological mechanism associated with a cell surface marker of the breast cancer stem cell population.

We further report the novel finding that p53 differentially regulates the expression of distinct target genes in different cell types based on its acetylation status, which is a function of Notch1 mediated repression of SIRT1. Specifically, our findings reveal a difference in p53 activity between differentiated epithelial cells and progenitor-like, tumor-initiating epithelial cells, which is recapitulated in breast cancer stem cells. The finding that iPS cells display similar regulation of the Notch1-mediated repression of SIRT1 suggests that it may be a functional marker of the differentiation status of cells. Of further significance is the fact that the p53-dependent events we observe occur in the absence of any exogenous cellular stress, i.e. genotoxic agents.

The results presented here underscore the previously observed tumor-initiating potential of BPE cells(12). In these cells loss of p53 results in the loss of a tumor suppressor, and the resultant upregulation of an oncogenic transcription factor, *c-myc*. In the case of differentiated epithelial cells, however, deletion of p53 does not result in the stimulation of *c-myc* expression. Therefore, differentiated epithelial cells possess a second barrier to transformation: the requirement to independently upregulate *c-myc* expression and/or activity. These results suggest that progenitor-like epithelial cells may be more amenable to transformation than differentiated epithelial cells and thus represent the cells of origin for human carcinomas. Finally, we demonstrate that regulation of p53 acetylation also has profound effects in cells harboring mutant p53. Specifically, acetylated mutant p53 stimulates the transcription of p73, which in turn mediates the chemoresistant property of breast cancer stem cells.

In summary, our results identify a novel Notch1 dependent mechanism of tumor initiation, cancer stem cell function and chemoresistance. By delineating the distinct signal transduction pathways and regulatory networks at work in tumor-initiating cells we may be able to develop therapeutics that target the earliest stages of tumor development, well before the malignant stage. Finally, our results suggest that while cancer stem cells may be intrinsically resistant to chemotherapy, they are also sensitive to cell killing by thrombospondin-1. Thus, therapies designed to mimic or stimulate Tsp-1 may have significant therapeutic efficacy against chemoresistant, triple negative breast cancer.

## Experimental Procedures

### Cell lines and constructs

The retroviral constructs pBabeZeo LT_c_ and pBabePuro LTK1 and pBabeHygro hTERT and the lentiviral shRNA vector pLK0.1 shp53 were a generous gift from Dr. Robert Weinberg, Whitehead Institute, Cambridge, MA. The p53 shRNA vector pMKO.1shp53-3UTR was a gift of Sean Downing, Foundation Medicine, Cambridge, MA and contained the following sequences: CATTCTGCAAGCACATCTG and was cloned into the Age1/EcoR1 sites in pMKO.1. The retroviral SV40 large T antigen mutant constructs, pBabePuro LT-Δ69-83, pBabeNeo LT_350_, pBabeNeo LT-D402H, and pBabeNeo LT-H42Q were a generous gift from Dr. James DeCaprio, Dana Farber Cancer Institute, Boston, MA. The lentiviral vectors specifying, Notch-sh1 and Notch-sh2 were purchased from Sigma-Aldrich and contained the following sequences: Notch-sh1: CCGGCCGGGACATCACGGATCATATCTCGAGATATGATCCGTGATGTCCCGGTTTTTG and Notch-sh2: CCGGCGCTGCCTGGACAAGATCAATCTCGAGATTGATCTTGTCCAGGCAGCGTTTTTG. The pYESir2 and PYESir2HY constructs were generous gift from Dr. Robert Weinberg. The pCMV-p53 and pCMV-p53R175H were generous gifts from Meredith Irwin, Hospital for Sick Kids, Toronto, On, CA. The lentiviral vector specifying shp63 and shp73 were designed with the following sequence: TGCCCAGACTCAATTTAGT, and GGATTCCAGCATGGACGTCTT. The Gateway Cloning vector pENTR223 TAP73 was a generous gift from Dr. Robert Weinberg, Whitehead Institute, Cambridge, MA.

The immortalization of human mammary epithelial cells (HME) was described previously (47). Human renal and lung (bronchial) epithelial cells (Lonza, Walkersville, MD) were immortalized via transduction of the retroviral vector pBabeHygro hTERT. Human dermal fibroblasts (hDF) were a generous gift from Dr. Robert Weinberg, and were immortalized via retroviral transduction with pBabeHygro hTERT to yield hFhT cells. HME, HRE, and hFhT cell lines expressing the SV40 large T antigen and LT mutants were generated by retroviral transduction with the pBabe vectors described above. Retroviruses were produced as previously described (47). Human dermal fibroblasts expressing pCMV and pCMV-E2F1 were produced through transient transfections as previously described (48). Human fibroblasts, HME cells and tumor initiating cells (BPE) expressing the lentiviral shRNA constructs described above were generated via lentiviral transduction as previously described (19, 49).

Both HME and HRE cell lines were cultured in a 1:1 mixture of F-12 Nutrient Mixture and Dulbecco’s Modified Eagle Medium (DMEM) (GIBCO, Carlsbad, CA) and supplemented with 5% fetal bovine serum (GIBCO), 1µg/ml hydrocortisone, 10ng/ml EGF, and 10μg/ml insulin (Sigma Chemicals, St. Louis, MO). Human bronchial epithelial cells were cultured in Bronchial Epithelial Growth Media (Lonza). Dermal and mammary fibroblasts were cultured in DMEM supplemented with 10% FBS. Lung fibroblasts were cultured in Minimum Essential Medium (GIBCO) supplemented with 10% FBS. HCT116, LNCaP, A549, MDA-MB-231, BT549, MCF-7 and MRC5 cell lines were obtained from American Tissue Type Collection (ATCC) and cultured in the prescribed media. SUM159 cell line was a generous gift from Dr. Stephen Ethier (University of South Carolina) and cultured in Ham’s F-12 (Life technologies) supplemented with 5% FBS, 5μg/ml Insulin and 1μg/ml Hydrocortisone. All cells were used for experimentation within 10 days of thawing.

iPS cells were a generous gift from Dr. George Daley, Boston Children’s Hospital, Boston, MA and were cultured in a 1:1 mixture of F-12 Nutrient Mixture and Dulbecco’s Modified Eagle Medium (DMEM) (GIBCO, Carlsbad, CA) and supplemented with 20% knockout serum replacer, 5mM L-Glutamine, 4ng/ml Basic FGF, 1% MEM-NEAA (Invitrogen, Carlsbad, CA) and 0.1mM 2-Mercaptoethaol (Sigma Chemicals).

### Western blotting

Cells were lysed in 50mM Tris-HCl (pH 7.5), 150mM NaCl, 1% Sodium deoxycholate, 1% NP-40, 0.1% SDS, 2nM DTT, and Complete Mini protease inhibitor cocktail (Roche, Mannheim, Germany). Western blots were performed as previously described using the following antibodies: *c-myc* (rabbit pAb, Cell Signaling Technologies, Danvers, MA), p53 (rabbit pAb, Cell Signaling Technologies), p21 (mAb clone DCS60, Cell Signaling Technologies), GAPDH (rabbit pAb, Trevigen, Gaithersburg, MD), SIRT1 (rabbit mAb E104, Abcam, Cambridge, MA), phosphorylation-p53 (Cell Signaling Technologies), Ac-p53 (rabbit pAb, Cell Signaling Technologies), Notch1 (rabbit mAb D1E11, Cell Signaling Technologies), β-actin (mouse mAb AC-15, Abcam), CD44 (Ab24504, Abcam), and MDM2 (PAB1729, Abnova, Taipei, Taiwan). Phosphorylation-Akt (S473) (mouse mAb D9W9U, Cell Signaling Technologies), Akt (rabbit mAb, Cell Signaling Technologies)

### Transient Transfections

Transient transfections were performed by adding 2-3μg of the appropriate vector in conjunction with 4-6μL Lipofectamine transfection reagent (Invitrogen). Cell culture media was changed 12 hours post-transfection and replaced with fresh media. Cells were harvested 48 hours post transfection and lysed for western blot analysis as described above.

### Drug treatment

For the p53 activity assay mammary epithelial cells were grown in the medium described above and dermal fibroblasts were grown in DMEM + 10% FBS. 500,000 cells were plated and cultured overnight, then medium was replaced and 50μM etoposide (EMD Millipore) was added to the medium. Cells were incubated with etoposide for 8 hours then were harvested and lysates were prepared for western blots (as described above). For drug treatment experiments, cells were incubated in culture medium for 1-16 hours with 25μM Nutlin-3 (Sigma), 25μM anacardic acid (EMD Millipore), 7.5μM trichostatin A (TSA) 20μM 6-Chloro-2,3,4,9-tetrahydro-1H-carbazole-1-carboxamide (SIRT1i) (EMD Millipore), 15μM SAHM1 (EMD Millipore), 25µM IMR1 (Sigma), 50-250μM rapamycin (EMD Millipore), 10μM LY294002 (EMD Millipore), 100-250nM Torin (EMD Millipore), 10μM MK-2206 (EMD Millipore) and 10μM MG132 (EMD Millipore). Cell lysates were then prepared and western blots performed as described above.

### Cell Proliferation Assays

For the cell proliferation assay all cells were grown in the same complete medium described above. 1,000 cells/well were plated in 96-well plates and incubated with 10μM Nutlin-3 with fresh Nutlin-3 added every 24h. The number of cells in culture was counted at 0h, 24h, 48h, 72h, 96h and 120h using WST-1 cell proliferation Reagent (Roche Diagnostics, Indianapolis, IN).

### Reverse-Transcriptase Real Time PCR

The expression level of c-myc, sirt1, TAp73, dNp73, TAp63, dNp63 were measured by real time quantitative PCR. Human glyceraldehyde phosphate dehydrogenase (GAPDH) was used as an internal control gene. The specific primers used are shown in Table 1. RNA was extracted from each sample cell line using Trizol reagent (Invitrogen). RNA concentration was measured using A_260_ in a Smartspec Plus spectrophotometer (Bio-Rad, Hercules, CA) and 1µg of RNA from each sample was converted to double stranded cDNA using iScript cDNA synthesis kit (Bio-Rad). RT-PCR was performed using iQ SYBR Green Supermix (Bio-Rad) on a MyiQ Single Color RT PCR Detection System (Bio-Rad). RT-PCR data was quantitated using the relative standard curve method and normalized against GAPDH intensity levels. P-values were calculated using one-way ANOVA.

### Chromatin Immunoprecipitation

Treated cells were cross-linked, lysed, immunoprecipitated, and DNA was extracted as per the protocol in the Actetyl-Histone H3 Immunoprecipitation (ChIP) assay kit (Millipore, Billerica, MA). Anti-acetyl-Histone H3 antibody was used for the immunoprecipitation (Millipore). Input samples were quantitated and RT-PCR was performed on the chromatin samples based on the concentrations of the input samples. Samples were assayed for the presence of the p53-binding site of the Myc promoter using the following primer set that amplifies a 208-bp fragment within the p53-response element of the human Myc gene: 5’-TGAGGGACCAAGGATGAGAAGAATG-3’ and 5’-TGAAAGTGCACTGTATGTAACCCGC-3’(50). Real-time (RT)-PCR data was quantitated using the relative standard curve method and was normalized against non-cross-linked controls. P-values were calculated using one-way ANOVA.

### Notch Inhibition

BPE cells were seeded one day before the treatment and synchronized in WIT-P medium (Cellaria, Cambridge, MA) containing 10 fold-diluted supplement. Synchronized cells were treated with Notch1 transcription factor inhibitor SAHM1 (15µM) (Millipore, Billerica, MA) for 24 hours, and gamma secretase inhibitor DAPT (25µM) (Millipore) for 16 hours. The cells were also treated with 20µM LY294002 (Millipore), 10µM MK2206 (Selleckchem, Houston, TX), 100nM, 250nM Torin (Millipore) and 100nM rapamycin (Millipore) for 1 hour and 16 hours respectively. Cells were then harvested and lysed in RIPA buffer containing protease inhibitor and phosphatase inhibitor. Cell lysate were subjected to immnoblotting as described above.

### FACS Analysis

Cells were collected and washed by PBS plus 2% FBS. Then cells were blocked by PBS plus 5% BSA and later incubated with FITC Anti-Human CD24 (BD Biosciences, San Jose, CA), and PE Anti-Human CD44 (BD Biosciences). Cells were washed 3 times each with PBS plus 2% FBS subsequent to incubation with primary antibodies. Finally cells were analyzed by BD FACSCalibur machine (BD Biosciences).

### Tumor sphere Formation Assay

Cells were collected when plates were around 60-70% confluent and resuspended into single cells in the tumor sphere medium. We plated 8,000 MDA-MB-231 and 10,000 BT-549 cells in 24-well ultra-low attachment plates (Corning Incorporated, Corning, NY). The sorted Notch high/ low cells were plated right after sorting at 1000 cells/ well. Tumor spheres were counted and measured at day 12. The tumor sphere medium was made as previously described and consisted of DMEM/F12, 100 U/ml penicillin/streptomycin, 20ng/ml EGF, 10ng/ml Basic FGF, 1 x B27 supplement.

### FACS Sorting

Cells were collected and washed by PBS plus 2% FBS. Then cells were blocked by PBS plus 5% Goat Serum (Vector Laboratories, Burlingame, CA) for 15 minutes at 4°C. Later cells were incubated with FITC Anti-Human CD24 (Abcam), V450 Anti-Human CD44 (BD Biosciences), PE Anti-Human EpCAM (BD biosciences) and Alexa Fluor 647 Anti-Human Notch1 (BD Pharmingen) for 30 minutes at 4°C in dark. Then Cells were washed 3 times with PBS plus 2% FBS. Finally cells were analyzed and sorted by BD FACSAria IIU machine (BD Biosciences). After sorting, cells were put into culture plates with full medium for 48 hours to recover, and then used for proliferation assay and *in vivo* tumor formation.

### Cancer stem cell Proliferation Assays

2,000 cells/well were plated in 96-well plates with the complete medium described above. The number of cells in culture was counted at 0h, 24h, 48h and 72h using WST-1 cell proliferation Reagent (Roche Diagnostics, Indianapolis, IN).

### Apoptosis Assay

FACS sorted cells were plated at 20,000 cell/ well in 12 well plate with complete medium. After 24 hours recovery, cells were treated with 20nM paclitaxel for 48 hours. At the end of the treatment, both of the floating cells and attached cells were collected for apoptotic analysis by Annexin V/ PI staining according to manufacturer’s protocol.

### *In vivo* tumor Formation

Notch1^hi^ and Notch1^low^ cells within the CD44^hi^/CD24^low^/ESA^+^ population of SUM159 breast cancer cells, were collected and resuspended in HBSS (Life technologies), then mixed with Matrigel 1:1 (BD Bioscience). The final cell concentration was 25 cells/μL. 500 cells in 20 μL were orthotopically injected into the mouse mammary fat pad and monitored by a Xenogen IVIS 200 system (Xenogen, Alameda, CA) once per week. The nicotinamide (Sigma Aldrich) treatment was initiated on the second day post-implantation at a dose of 150 mg/kg/day. At the end of the experiment, mice were sacrificed and mammary fat pad were collected for future assay.

### Patient TMA Notch1 staining

We used a population based series of 534 breast carcinomas from the period 1996 - 2003, as previously described (51). The patients received treatment according to standard protocols in a single institution. Follow-up information was given by the Norwegian Cause of Death Registry, and can be considered accurate and complete. Last date of follow-up was December 31, 2011. Outcome data include survival status, survival time and cause of death. During the follow-up period, 79 patients (15%) died from breast carcinoma, and 62 (12%) died from other causes. The median follow-up time was 13 years estimated by the reverse Kaplan-Meier method. The study was approved by the Western Regional Committee for Medical and Health Research Ethics (Norway), REC West (REK 2012/1704).

### Clinico-pathological variables

The following variables were recorded: age at diagnosis, tumor diameter, histologic type, histologic grade, lymph node status, and hormonal receptor status. Furthermore, the results of previously investigated biomarkers (HER2 and Ki67) were included(51).

### Notch 1 immunostaining

Immunohistochemistry was performed on 5µm standard sections of formalin fixed, paraffin embedded tumor tissue microarrays. De-waxing with xylene/ethanol at different concentrations (100%, 96% and then 80%) followed by about 20 minutes heat-induced targeted epitope retrieval using PH6, DAKO S1699 in microwave. The sections were then allowed to cool down about 20 minutes. An endogenous enzyme block was used to block the endogenous enzymatic activity (DAKO K4007). The slides were then incubated with primary polyclonal rabbit antibody; Notch 1(Abcam ab8925) at 1:200 dilution for 60 minutes. The sections were then incubated with the secondary antibody; Envision system-HRP (DAKO K4007) in 30 minutes. The visualization was performed by diaminobenzidine (DAB) as chromogen for 10 minutes, followed by hematoxylin and eosine for 3 minutes.

### Staining index

Staining was assessed by a semi-quantitative and subjective grading system that considers intensity of staining and proportion of tumor cells showing positive staining. For Notch 1, a staining index (values = 0–9) was determined by multiplying the score for intensity of staining (none = 0, weak = 1, moderate = 2 and strong = 3) with the score for proportion of tumor cells stained (<10% = 1, 10%–50% = 2, >50% = 3).

## Acknowledgements

We would like to thank George Daley, Robert Weinberg, Meredith Irwin, and Sean Downing for reagents and Kristin Johnson for artwork. We also would like to thank Judah Folkman, Richard O. Hynes, Bruce Zetter, Philip Iaquinta and Sarah Huntwork for helpful discussions. RW was supported by NIH R01 CA135417, the Elsa U. Pardee Foundation and the Strike 3 Foundation.

