## Supplementary Figures for "Notch1 regulates breast cancer stem cell function via a non-canonical cleavage-independent pathway"

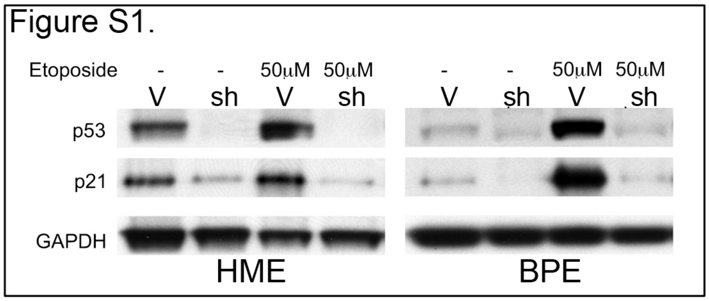


**Supplementary Figure 1. Effects of silencing p53 on p21 induction**
Immunoblot analysis of p53, p21 and GAPDH expression in human mammary epithelial cells (HME) and human tumor initiating mammary epithelial cells (BPE) transduced with empty pLKO.1 vector (V) or pLKO-shp53 (sh), that were untreated (-) or treated with 50μm etoposide.

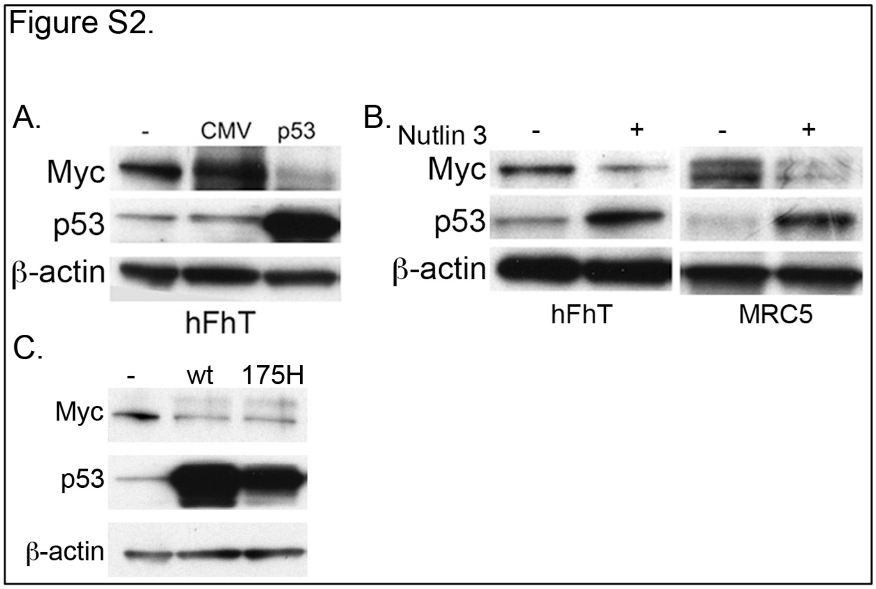


**Supplementary Figure 2. Effects of p53 modulation on Myc expression in fibroblasts**
Immunoblot analysis of:
**(A)** Myc, p53 and actin expression in telomerase immortalized human foreskin fibroblasts that were untransfected or transfected with empty pCMV vector, or pCMV-p53;
**(B)** Myc, p53 and actin expression in telomerase immortalized human foreskin fibroblasts and normal MRC5 lung fibroblasts that were untreated (-) or treated with 25μm Nutlin-3;
**(C)** Myc, p53 and actin expression in telomerase immortalized human foreskin fibroblasts that were untransfected or transfected with pCMV-p53, or pCMV-p53R175H.

Figure S3.

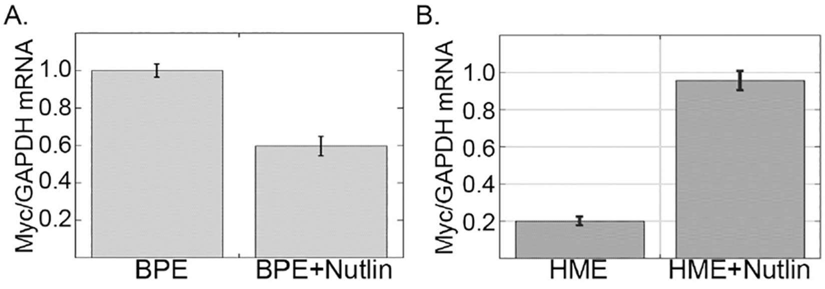


**Supplementary Figure 3. Differential regulation of the myc promoter by p53 in different cell types.**
(A) Relative expression of Myc mRNA, normalized to GAPDH, in BPE cells that were untreated (-) or treated with Nutlin-3;
(B) Relative expression of Myc mRNA, normalized to GAPDH, in HME cells that were untreated (-) or treated with Nutlin-3;

Figure S4.

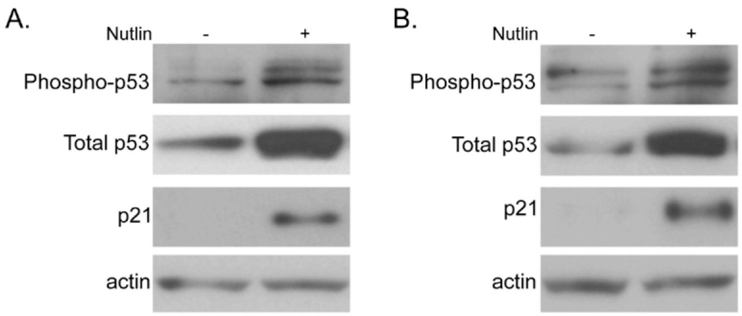


**Supplementary Figure 3. Effects of Nutlin-3 treatment on p53 phosphorylation**
Immunoblot analysis of phospho S15,S21 p53, total p53, p21 and actin expression in **(A)** BPE cells and **(B)** HME cells that were untreated (-) or treated with 25μm Nutlin-3.

Figure S5.

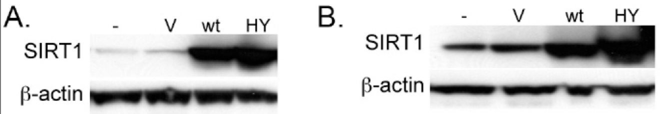


**Supplementary Figure 4. Overexpression of wt and mutant SIRT1**
Immunoblot analysis of:
**(A)** SIRT1 and actin expression in BPE cells that were untreated (-) or transduced with pBABEpuro (V), pYE-SIRT1 (wt) pYE-SIRT1HY (HY);
**(B)** SIRT1 and actin expression in HME cells that were untreated (-) or transduced with pBABEpuro (V), pBABEpuro-SIRT1 (wt) pBABEpuro-SIRT1HY (HY);

Figure S6.

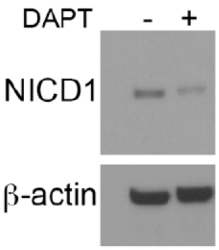


**Supplementary Figure 5. Effects of DAPT on NICD1 production.**
Western blot of NICD1 and β-actin expression in BPE cells that were untreated (-) or treated with 25μM DAPT.

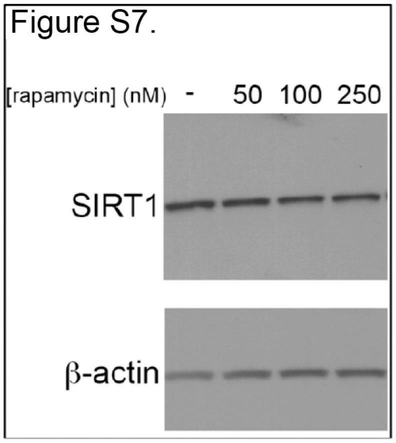


**Supplementary Figure 7. Effects of Rapamycin on SIRT1 expression.**
Western blot of SIRT1 and β-actin expression in BPE cells that were untreated (-) or treated with 50, 100 and 250nM rapamycin.

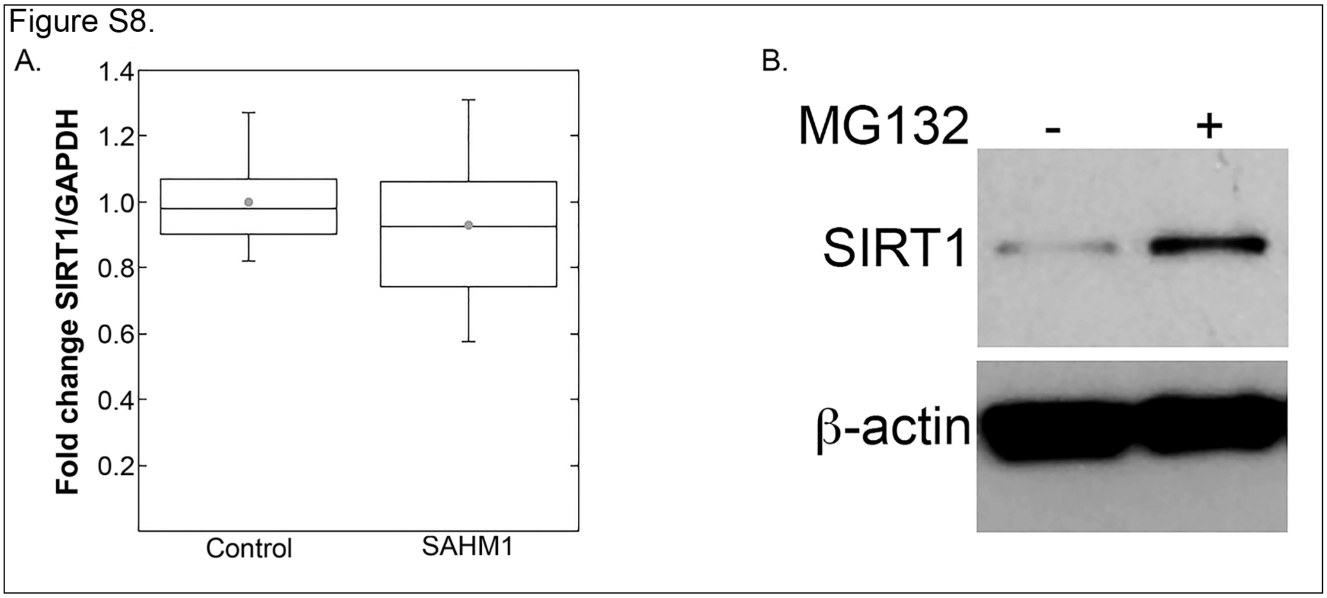


**Supplementary Figure 8. Notch1 downregulates SIRT1 via proteasomal degradation.**
**(A)** Box plot of fold change of SIRT1 mRNA levels as determined by RT-PCR;
**(B)** Western blot of SIRT1 and β-actin expression in BPE cells that were untreated or treated with the proteasome inhibitor MG132 overnight.

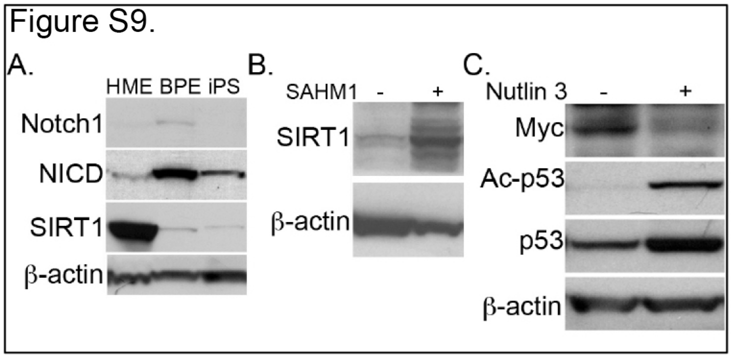


**Supplementary Figure 9. Analysis of Notch1 and p53 activity in iPS cells**
Immunoblot analysis of:
**(A)** Notch, SIRT1 and actin expression in BPE, HME and human iPS cells;
**(B)** SIRT1 and actin expression in human iPS cells that were untreated (-) or treated with SAHM1 (+) for 24 hours;
**(C)** Myc, Ac-p53, p53 and actin expression in human iPS cells that were untreated (-) or treated with 25mM Nutlin-3 (+) for 8 hours.

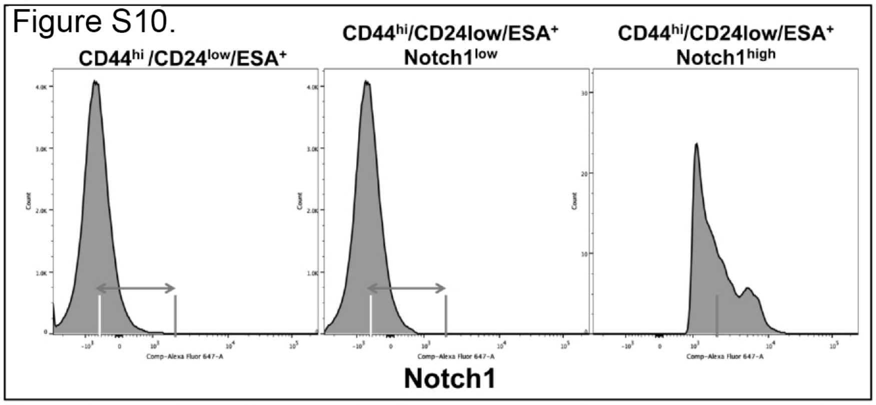


**Supplementary Figure 10. Analysis of Notch1 expression in different populations of MDA-MB-231 cells**

Histogram of FACS analysis of Notch1 expression in $CD44^{hi}/CD24^{low}/ESA^{+}$, $CD44^{hi}/CD24^{low}/ESA^{+}/Notch1^{low}$, and $CD44^{hi}/CD24^{low}/ESA^{+}/Notch1^{high}$ cells isolated from the bulk population of MDA-MB-231 cells by FACS sorting.

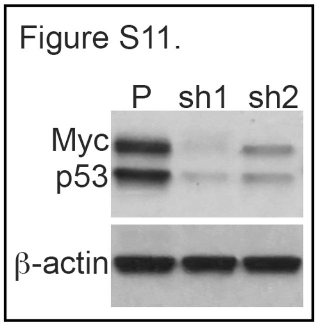


**Supplementary Figure 11. Silencing of p53 via two different shRNA sequences.**

Western blot of Myc, p53 and β-actin in SUM159 cells following lentiviral transduction of two independent shRNA sequences targeting the coding region (sh1) and 5’ UTR (sh2) of the gene.

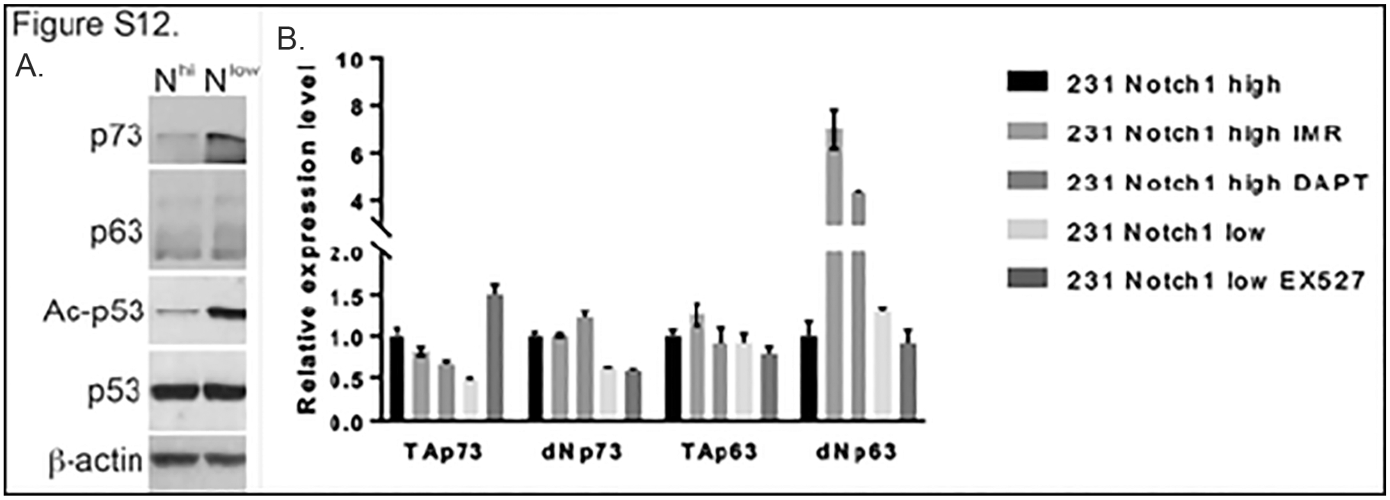


**Supplementary Figure 12. Analsys if p53 family member expression in MDA-MB-231 cells.**

Western blot of p73, p63, acetylated p53( Ac-p53), total p53, and β-actin in Notch1 high and low expressing MDA-MB-231 cells. **(B)** RT-PCR of TA and DN isoforms of p73 and p63 in Notch1 high MDA-MB-231 cells hat were untreated or treated with IMR1, DAPT, and low Notch1 expressing MDA-MB-21 cells that were untreated or treated with EX527.
